# SPLICER: A Highly Efficient Base Editing Toolbox That Enables *In Vivo* Therapeutic Exon Skipping

**DOI:** 10.1101/2024.04.01.587650

**Authors:** Angelo Miskalis, Shraddha Shirguppe, Jackson Winter, Gianna Elias, Devyani Swami, Ananthan Nambiar, Michelle Stilger, Wendy S. Woods, Nicholas Gosstola, Michael Gapinske, Alejandra Zeballos, Hayden Moore, Sergei Maslov, Thomas Gaj, Pablo Perez-Pinera

## Abstract

Exon skipping technologies enable exclusion of targeted exons from mature mRNA transcripts, which has broad applications in molecular biology, medicine, and biotechnology. Existing exon skipping techniques include antisense oligonucleotides, targetable nucleases, and base editors, which, while effective for specific applications at some target exons, remain hindered by shortcomings, including transient effects for oligonucleotides, genotoxicity for nucleases and inconsistent exon skipping for base editors. To overcome these limitations, we created SPLICER, a toolbox of next-generation base editors consisting of near-PAMless Cas9 nickase variants fused to adenosine or cytosine deaminases for the simultaneous editing of splice acceptor (SA) and splice donor (SD) sequences. Synchronized SA and SD editing with SPLICER improves exon skipping, reduces aberrant outcomes, including cryptic splicing and intron retention, and enables skipping of exons refractory to single splice-site editing. To demonstrate the therapeutic potential of SPLICER, we targeted *APP* exon 17, which encodes the amino acid residues that are cleaved to form the Aβ plaques in Alzheimer’s disease. SPLICER reduced the formation of Aβ42 peptides *in vitro* and enabled efficient exon skipping in a mouse model of Alzheimer’s disease. Overall, SPLICER is a widely applicable and efficient toolbox for exon skipping with broad therapeutic applications.

## Introduction

Programmable genome-editing nucleases have fundamentally transformed biotechnology and medicine by providing a facile means to introduce targeted modifications in the genome of living cells.^1^ Nonetheless, their reliance on DNA double-strand breaks (DSBs) has been linked to undesirable effects, including chromosomal deletions^2^, chromotripsis^3^, activation of the p53-mediated damage response pathway^4^, and mRNA misregulation^5^, all of which can hinder their implementation.^6,7^ For these reasons, emergent gene editing technologies capable of introducing precise and predictable mutations without DSBs are becoming increasingly adopted for a range of applications.

One such technology are base editors (BEs), which utilize a fusion of a Cas9 nickase and a cytosine deaminase to introduce C>T (cytosine base editors (CBEs)) or an adenosine deaminase (adenosine base editors (ABEs)) to introduce A>G mutations at genomic loci with complementarity to a single-guide RNA (sgRNA)^8,9^. One application of BEs is the programmed modulation of RNA splicing, which can be used to modify the transcriptional landscape of a cell for basic biology and therapeutic applications. For this goal, we previously described CRISPR-SKIP, a technology that relies on BEs to target and mutate highly conserved AG dinucleotide motifs in splice acceptor (SA) sequences, which can prevent their recognition by the spliceosome and induce skipping of the corresponding exon.^10^ In addition to our work, others have utilized SpCas9 to introduce exon skipping by mutating splice donor (SD) sites and have applied BEs to induce exon skipping for various applications.^11–15^ Notably, by excluding exons from mature mRNA transcripts, CRISPR-SKIP and other approaches for exon skipping hold the potential to eliminate mutated sequences and recover reading frames lost to chromosomal deletions.

Nonetheless, multiple challenges exist that prevent the broad implementation of SpCas9-comprised BEs for exon skipping. First, the native SpCas9 protein can only target splice sites with NGG protospacer adjacent (PAM) motifs nearby the target locus^1,16^, a restriction that can limit the number of exons that can be effectively engaged. Second, the adenosine and cytosine deaminase domains most commonly used in first-generation BEs possess context-dependent preferences that can prevent their effective editing of a range of splice sites.^1,17,18^ Third, even when SpCas9 BE technologies are able to edit splice sites, there exist in many exons cryptic splice sites that can be recognized by the spliceosome, which can lead to partial skipping.^13,19,20^ Further, in addition to cryptic splicing sequences, the targeting of SA or splice donor (SD) sites can lead to the partial or full retention of introns in mature transcripts, which can result in the incorporation of novel sequences into open reading frames or create early termination codons that could undesirably repress expression.^19–23^

To overcome these limitations, we develop SPLICER, a toolbox of near-PAMless BEs comprised of enhanced deaminase domains that enable the simultaneous targeting of SD and SA sites, which we show improves exon skipping, reduces aberrant splicing outcomes, including cryptic splicing and intron retention, and enables the skipping of exons previously found to be resilient to skipping. Further, to highlight the potential of this toolbox, we demonstrate that SPLICER can be used to induce skipping of exon 17 in the gene encoding the amyloid precursor protein (APP), a transmembrane receptor that contributes to the formation of the amyloid deposits in the brain of patients afflicted by Alzheimer’s disease.^24–30^. SPLICER reduced the formation of Aβ42 peptides *in vitro* and mediated efficient exon skipping in a mouse model of Alzheimer’s disease.

Given its precision, versatility and efficacy, SPLICER will have broad applications in medicine and biotechnology.

## RESULTS

### Engineered near-PAMless SpCas9 variants enable targeting of SAs inaccessible by the native SpCas9

One limitation of SpCas9 for exon skipping is its reliance on the NGG PAM motif. We previously demonstrated that the use of SaCas9 or SpCas9-VQR could alleviate this problem, but BEs comprised of these variants often exhibit lower activity than those composed of the wild-type (WT) variant.^10,31^ Recently, several versions of SpCas9 with relaxed PAM preferences have been developed, including SpCas9-NG (NG PAM)^32^, xCas9-NG (NG PAM)^33,34^, NAG-Cas9 (NRG PAM)^35^, SpCas9-NRNH^36^ (NRNH PAM), SpG Cas9 (NRN PAM)^37^, and SpRY Cas9 (near NNN

PAMs).^37^ Most notably, SpRY Cas9 has demonstrated DNA editing activity at NRN and NYN PAM sites comparable to SpCas9 when targeted to its native NGG PAM.^37^ SpRY Cas9 was also demonstrated to edit a larger number of compatible targets compared to xCas9-NG, SpCas9-NG, and SpG Cas9.^37^ Importantly, we observed that SpRY Cas9 could install base edits using NDN PAMs, which is particularly convenient for targeting SAs, whose consensus sequence is A/T-rich and would enable tiling of splicing sequences with multiple sgRNAs (**Fig. 1A**).

**Figure 1.**
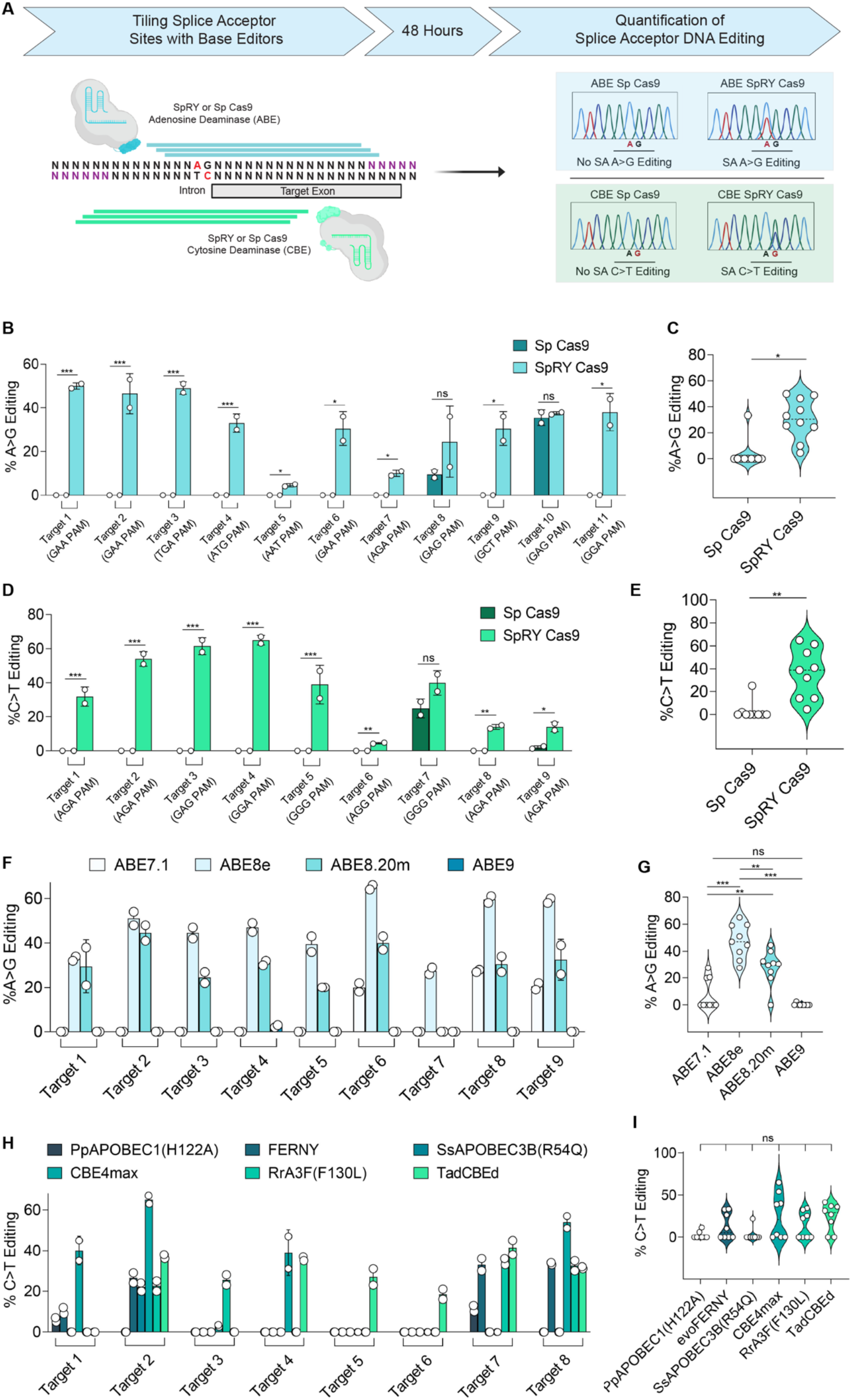
Spry Cas9 in Combination with Different Deaminases Enables Efficient Disruption of Targeted Splice Acceptors. **(A)** Schematic representation of the overall approach to disrupt splicing elements with near-PAMless base editors (BEs) by tiling sgRNAs. **(B-E)** Comparison of SpCas9 and SpRY Cas9 editing efficiency at the splice acceptors (SAs) of eleven target exons using adenosine base editors (ABEs) **(B, C)** or cytosine base editors (CBEs) **(D, E)**. Summary of editing efficiency at individual targets accomplished with Sp Cas9 or SpRY Cas9 ABEs (**C**) or CBEs (**E**). Comparison of DNA editing efficiency at multiple SAs using SpRY Cas9 fused with one of four different adenosine deaminases **(F, G)** or one of six different cytosine deaminases **(H, I)**. Summary of editing efficiency at individual targets accomplished with different SpRY ABEs (**G**) or different SpRY CBEs (**I**). n = 2; ns, no significance; *, p<0.05, **; p<0.01; *** p<0.001; two-tailed, unpaired t-test except **G** and **H**, which were analyzed via One-Way ANOVA with Tukey’s Post-hoc.

Given the broader targeting range observed with SpRY Cas9, we next tested the exon skipping ability of a BE toolbox comprised of this variant. To accomplish this, we targeted multiple exons by tiling SAs with eight sgRNAs per gene at six exons that contained either one NGG PAM or no NGG PAMs with the adenosine deaminase ABE8e^38^ (**Fig. 1B, C**) or the cytosine deaminase CBE4max (**Fig. 1D, E**) with either SpCas9 or SpRY Cas9 in HEK293T cells. Using ABE8e and APOBEC1-CBE4max, we identified at least one sgRNA that introduced the desired mutation for every target exon for SpRY Cas9, while BEs based on SpCas9 only introduced mutations in the SA of two exons (**Fig. 1B-E**). At target SAs with NGG PAMs both SpCas9 and SpRY Cas9 introduced the target change at similar rates (p > 0.05, **Fig. 1B, D)**. Overall, the mean editing rates for SpRY Cas9-ABE8e and SpRY Cas9-CBE4max were ∼31% and ∼35%, respectively, while the mean editing rates for WT ABE8e and CBE4max were ∼3% for each. Thus, SpRY Cas9 provided a 9.5- and 9.7-fold increase in mean SA editing when comparing WT to SpRY for both ABE8e and CBE4max, respectively, (p < 0.05; **Fig. 1C, E**).

An important parameter for optimizing editing of a splicing sequence by a BE is the selection of the deaminase domains, as multiple new Cas9 BEs have been developed for various purposes. For example, in addition to CBE4max^39^, CBEs composed of the evoFERNY^40^, SsAPOBEC3B (R54Q)^41^, RrA3F (F130L)^41^, TadCBE^42^, and PpAPOBEC1 (H122A)^41^ catalytic domains have expanded editing contexts, increasing overall editing efficiency, and decreasing non-specific deamination. In the case of ABEs, evolved variants such as the ABE8^43^ family, including ABE8e^38^ and ABE8.20m^43^ have increased editing efficiency and different editing windows than the previous generation ABE7.10.^43^ Additionally, ABE9^44^ was developed to minimize bystander editing by BEs of the ABE8 family. When fusing different deaminases to SpRY Cas9, we observed distinct profiles of edited SA sequences (**Fig. 1F-I**).

When comparing ABEs, ABE8e outperformed all other deaminases possessing a mean editing rate of ∼47% compared to ∼7%, ∼28%, and ∼0.3% for ABE7.1, ABE8.20m, and ABE9, respectively (p < 0.01, **Fig. 1F, G**), which correlates well with previous studies demonstrating the superior editing activities of ABE8e^38^ and ABE8.20m.^43^

Comparisons of CBEs revealed different editing profiles, but we did not observe any significant difference in mean editing rates among the groups (p > 0.05, **Fig. 1H, I**). The most efficient CBEs were TadCBEd, which successfully edited 75% of the targeted SAs (mean editing rate ∼32%) followed by CBE4max (50%, mean editing rate ∼40%), evoFERNY (50%, mean editing rate ∼26%) and RrA3F (F130L) (50%, mean editing rate ∼29%). SsAPOBEC3B (R54Q) and PpAPOBEC1 (H122A) only edited 1 and 2 target sites respectively at rates lower than ∼22% (**Fig. 1H, I**). Interestingly, TadCBE and RrA3F (F130L) enable editing of target sequences that were resilient to modification by every other BE system tested (**Fig. 1I**).

Thus, fusion of SpRY Cas9 with different deaminases significantly increased the number of exons that can be targeted with high efficiency.

### Simultaneous editing of SA and SD sites enhances exon-skipping

While there are different parameters that can influence exon splicing, a simplistic model of exon skipping would suggest that exon skipping rates are the result of the equilibrium between the DNA editing rate that disrupts different splicing elements favoring skipping and the persistence of non-edited functional splicing elements that favors exon inclusion, such as splicing enhancers or cryptic splicing elements. Thus, we reasoned that exon skipping could be enhanced by simultaneously targeting multiple splicing elements, such as SA and SD sites (**Fig. 2A**), could shift the equilibrium further towards exon skipping.

**Figure 2.**
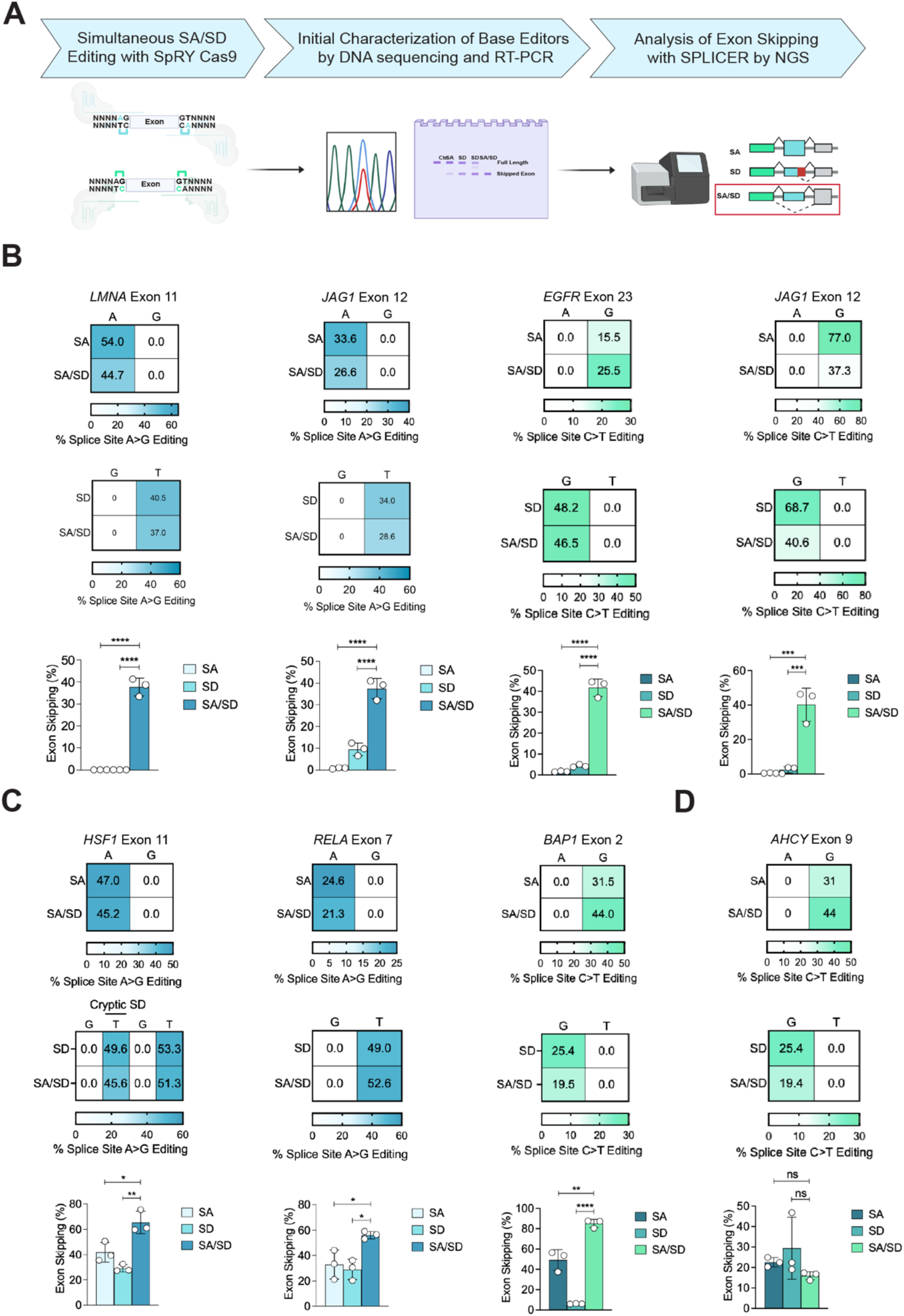
Simultaneous Editing of Splice Sites with SPLICER Enhances Exon Skipping. **(A)** Illustration of SPLICER strategy for simultaneously targeting SAs and SDs with ABEs or CBEs followed by screening to identify effective BE systems and characterization by next generation sequencing (NGS). **(B-D)** Editing of AG dinucleotides in the SA (top panel) and GT dinucleotides in the SD (medium panel) and exon skipping (bottom panel) with ABEs (highlighted blue) or CBEs (highlighted green). Improvements in exon skipping were synergistic when targeting *LMNA* exon 11, *JAG1* exon 10, *EGFR* exon 23 and *JAG1* exon 12 (**B**). Improvements in exon skipping were additive when targeting *HSF1* exon 11 and *RELA* exon 7 (**C**). Editing of SA and SD of *AHCY* exon 9 with ABEs did not improve exon skipping rates compared with editing of SA or SD alone (**D**). n=3 for all experiments; ns, no significance; *, p<0.05; ** p<0.01; **** p<0.0001; One-Way ANOVA, Tukey’s Post Hoc comparing SA/SD to SA and SD.

To test this hypothesis, we targeted the SDs and SAs, referred to as dual splice site targeting, of 4 exons with SpRY Cas9-ABE8e or SpRY Cas9-CBE4max and performed high-throughput DNA sequencing to analyze splicing induced by selected sgRNA pairs (**Fig. 2B, C, D**). At steady state, we did not detect alternative splicing events at any of the targeted exons in control samples (**Fig. S1A**). Following treatment with ABEs, we observed that dual targeting improved exon skipping rates across all targets for multiple sets of sgRNAs (**Fig. 2B, Fig. S1B, S1C)**, with effects ranging from synergistic (**Fig. 2B**) to additive (**Fig. 2C**) in comparison with targeting just the SA or just the SD. For example, when targeting *HSF1*, exon skipping improved from 40%, which was achieved by targeting the SD alone, to 58% when both splice sites were targeted (p = 0.013, **Fig. 2C)**. When targeting *RELA* exon 7 the improvement in exon skipping was additive, increasing from 32.6% and 25.3% skipping when targeting the individual SA and SD sites, respectively, to 56% when targeted simultaneously (SA/SD vs SA, p = 0.031, SA/SD vs SD, p = 0.014). Moreover, in other scenarios such as targeting exon 12 of *JAG1*, exon skipping improved from 0% and 9% when targeting the individual SA or SD sites, respectively, to 37% when both sites were targeted simultaneously (SA/SD vs SA, p < 0.0001, SA/SD vs SD, p < 0.0001, **Fig. 2B**), enhancing skipping synergistically. In some cases, exon skipping was accomplished despite limited skipping when targeting the individual single splicing sequences, such as for *LMNA* exon 11, where skipping increased from undetectable to 37% when editing both splice sites (SA/SD vs SA, p = 0.0001, SA/SD vs SD, p < 0.0001, **Fig. 2B**).

Dual splice site targeting of SAs and SDs with CBEs improved skipping in 75% of all targeted exons, with the improvements found to vary in a similar manner to the ABEs. Like for the ABEs, additive and synergistic trends in exon skipping were observed for multiple sets of sgRNAs targeting a specific exon (**Fig. S1B, S1C**). For example, the increase in exon skipping was additive for *BAP1* exon 2 (SA/SD vs SA, p = 0.0011, SA/SD vs SD, p < 0.0001, **Fig. 2C**), while exon skipping with CBEs was synergistic for both *EGFR* exon 23 (SA/SD vs SA, p < 0.0001, SA/SD vs SD, p < 0.0001) and *JAG1* exon 12 (SA/SD vs SA, p = 0.0003, SA/SD vs SD, p = 0.0004, **Fig. 2B**). Interestingly, when targeting *JAG1* exon 12, exon skipping increased ∼10-fold despite a decreased in DNA editing rates at the SA and SD sites by ∼50% when simultaneously targeting both the SA and SD (**Fig. 2B**), which highlights the large impact that dual targeting of splice sites can have on exon skipping even when only relative low DNA editing is accomplished. Finally, for *AHCY* exon 9, simultaneous editing did not improve exon skipping which was ∼28% (SA/SD vs SA, p = 0.654, SA/SD vs SD, p = 0.233, **Fig. 2D**), despite editing the SA and SD sites at efficiencies of ∼44% and ∼20% respectively.

Except for targeting of *JAG1* exon 12 with CBEs, when comparing single and dual splice site editing scenarios, genomic DNA editing at the splice sites was not significantly different (p > 0.05), indicating that the improvement in exon skipping was not due to higher editing rates but likely the effect of disrupting both SAs and SDs.

These results highlight that simultaneous targeting of multiple splicing sequences with SPLICER enhances full exon skipping outcomes. Further, they also suggest that one mechanism for the improvement of exon skipping rates is the suppression of aberrant splicing outcomes.

### SPLICER lowers cryptic splicing and intron retention

One problem frequently observed when inducing exon skipping is the presence of splicing aberrations, such as cryptic splicing and intron retention. Cryptic splicing occurs when the spliceosome machinery utilizes sites that closely resemble the canonical SA or SD sequences, resulting in a transcript in which the target exon is not fully skipped.^13,45,46^ Similarly, intron retention can occur when disruption of a SA or SD does not induce full exon skipping; instead, part of an intron or the entire intron is incorporated into the mature mRNA.^13,22,47^ While events such as these are complex and not fully understood, we and others^13,47–49^ have observed that ‘AG’ and ‘GT’ bases near the consensus SD and SAs are frequently interpreted by the spliceosome machinery as cryptic SAs or SDs respectively (**Fig. 3**), which can prevent full skipping of an exon when only one splicing element is disrupted.

**Figure 3.**
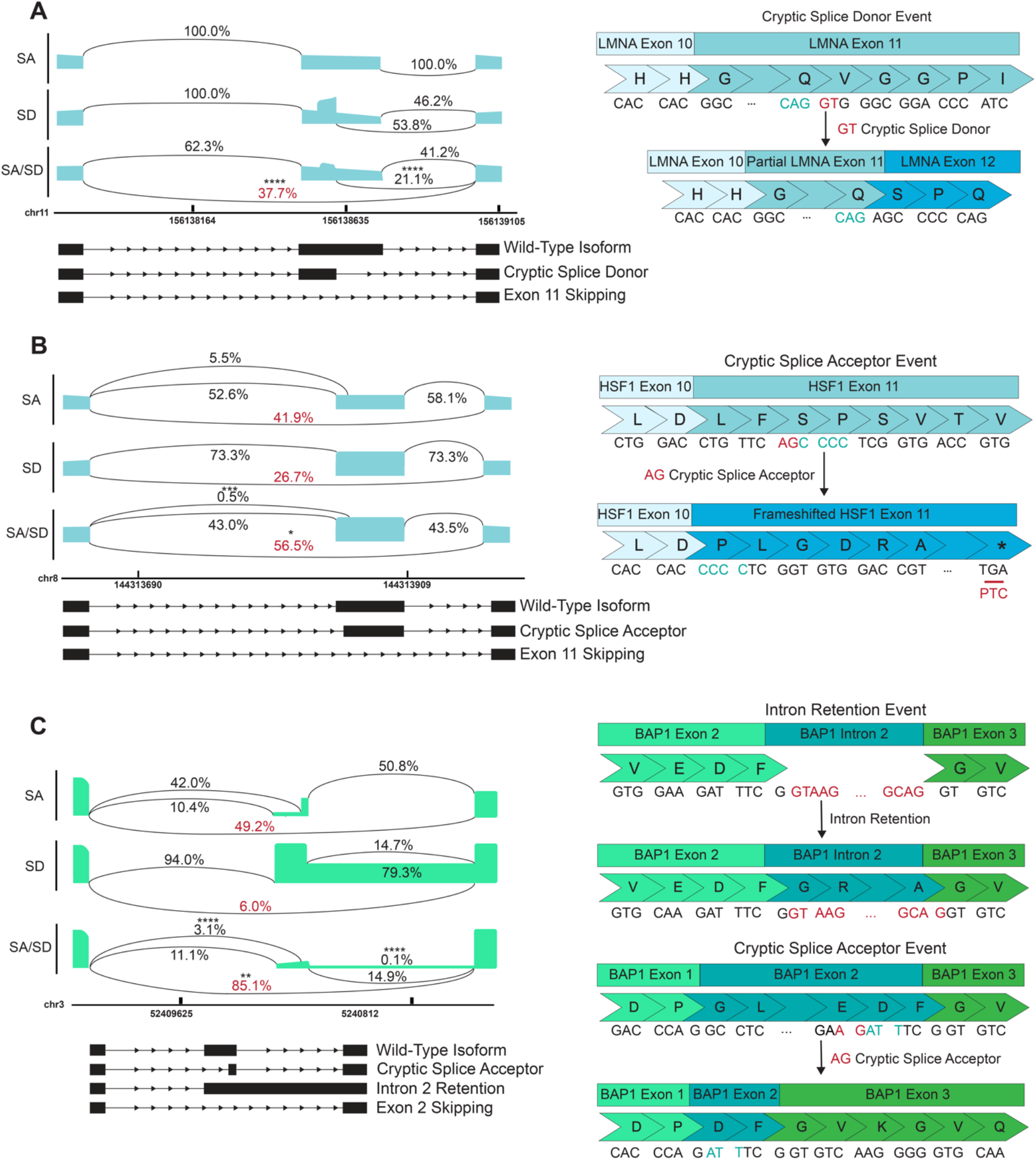
SPLICER Improves Exon Skipping Through Reduction of Both Cryptic Splicing and Intron Retention. (**A)** Sashimi plot describing exon splicing following ABE editing of *LMNA* exon 11 with SPLICER, in which a cryptic SD site is recognized in the middle of the exon when editing the SD (left), leading to no exon skipping and partial retention of exon 11 (right). This aberrant splicing event is reduced 3-fold by ABE editing of both SA and SD sites. **(B)** Sashimi plot representing splicing of *HSF1* exon 11 before and after ABE editing of splicing elements demonstrating a cryptic SA recognized in *HSF1* exon 11 (left). This cryptic splicing leads to a frameshift mutation in a protein, which results in a premature termination codon in exon 11 (right). Cryptic splicing is reduced 10-fold when editing both sites (left). **(C)** Sashimi plot describing splicing of *BAP1* exon 2 following editing with CBEs (left). Cryptic splicing and intron retention occur with SA and SD editing alone, leaving an in-frame, but mutated protein (right). Both events are minimized with SPLICER (left). n=3; *, p<0.05; **, p<0.01; ***, p<0.001; ****, p<0.0001; One-Way ANOVA, Tukey’s Post Hoc comparing cryptic splicing events in SA or SD to the rate of that cryptic splicing event in simultaneous SA/SD editing.

Since editing of both SA and SD sites increased full exon skipping, we hypothesized that one reason for the increase in full-length exon splicing could be a decrease in the rate of cryptic splicing. To test this hypothesis, we targeted the SA, the SD, or both in exon 11 of *LMNA*, where editing a single splicing sequence has been shown to induce cryptic splicing (**Fig. 3A**). While targeting the SA of *LMNA* exon 11 failed to induce exon skipping, targeting the SD led to ∼53% partial exon 11 skipping, where a ‘GT’ dinucleotide that is normally part of the codon corresponding to residue V607 can be recognized as a cryptic SD (**Fig. 3A**), leading to partial exon skipping and a new reading frame for *LMNA*. However, the rate of full exon 11 skipping when targeting both the SD and SA was ∼37%, while the rate of partial exon skipping was found to decrease ∼3-fold (SA/SD vs SD, p = 0.0001). This change in cryptic splicing was also observed at the SA for targets such as *HSF1* exon 11 (**Fig. 3B**). In this instance, a cryptic ‘AG’ sequence present at S419 can be utilized, leading to a frameshifted transcript with a premature termination codon (PTC) within exon 11. We found that this cryptic splicing event was reduced ∼10-fold upon simultaneous splice site disruption (SA/SD vs SA, p = 0.0005). Additionally, in *HSF1* exon 11, there exists a cryptic SD created by a ‘GT’ motif located at the 5’ of the canonical SD, which can be used as a splice-site, leading to a frameshift that extends through exons 12 and 13 and can result in a stop codon after the canonical stop codon in exon 13. When targeting the SD of *HSF1* exon 11, the cryptic SD event was recognized in ∼4.8% of cases. Further, this cryptic event was reduced by ∼4.9-fold to 0.97% for dual splice site editing (p < 0.0001, **Fig. S2A**). Similarly, when targeting the SA of *BAP1* exon 2, the full exon skipping rate was measured to be ∼49%, while ∼42% of the transcripts were found to contain a partial exon 2 resulting from a cryptic SA (**Fig. 3C**). This cryptic splicing event leads to a transcript containing a truncated exon 2, although the reading frame was preserved. When targeting the SD, full exon skipping was only detected in ∼6% of the transcripts, while ∼79% of the transcripts retained intron 2 fully. Both aberrations were minimized by targeting both SD and SA sequences, which decreased cryptic splicing to ∼3% (SA/SD vs SA, p = 0.0003,), reduced intron retention to <1% (SA/SD vs SD, p < 0.0001), and provided ∼85% full length exon skipping. These results support that SPLICER improves full exon skipping by dual splice site DNA editing through the reduction of cryptic splicing events.

### SPLICER reduces in vitro Aβ42 expression and skips APP exon 17 in vivo

Given the efficiency of exon skipping accomplished with SPLICER, we next sought to determine whether it could be utilized to skip exons with therapeutic value, such as exon 17 in the *APP* gene which, in our experience, was resilient to skipping with CRISPR-SKIP. *APP* is located in chromosome 21 and encodes the amyloid precursor protein (**Fig. 4A**).^24,25^ APP digestion by β- and γ-secretase generates the Aβ42 peptide, which is hypothesized to contribute to the pathogenesis of Alzheimer’s disease (AD) (**Fig. S3A**).^24,50^ Importantly, this cleavage site is encoded within exon 17, an exon whose skipping has been previously shown to reduce the formation of Aβ42 plaques^25,51^. While previous attempts at editing the SA of *APP* exon 17 with SpCas9-derived BEs had failed in our laboratory, we found that the PAM flexibility afforded by SpRY Cas9 enabled efficient editing of the SA with both ABE8e (50%) and a CBE4max (16%) in HEK293T cells (**Fig. 4B-C**). We also utilized SpRY Cas9 to target the SD of *APP* exon 17, identifying three sgRNAs that edited it at efficiencies of 78%, 67%, and 16% (**Fig. 4C-D**).

**Figure 4.**
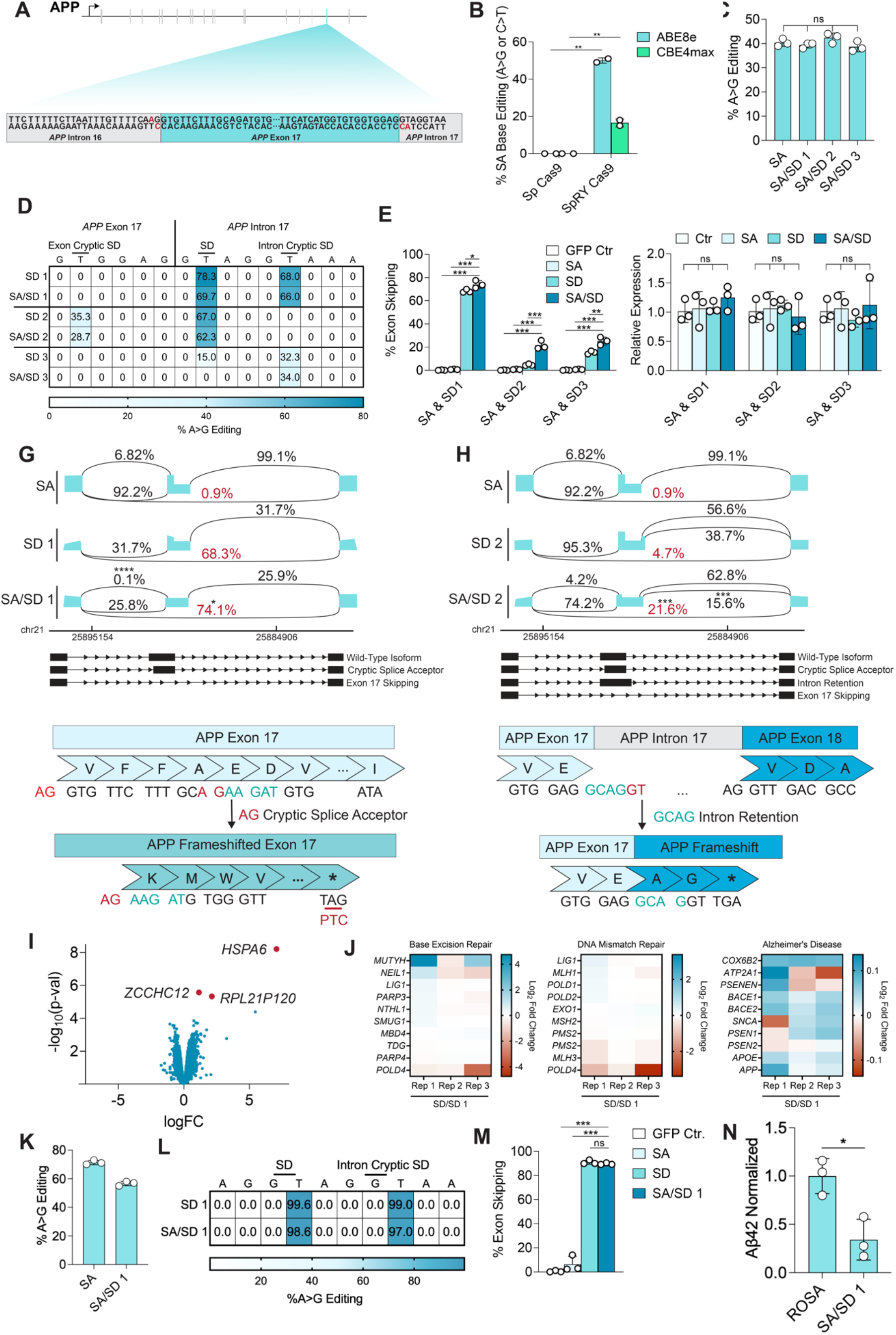
SPLICER Skips APP Exon 17 *In Vitro* And Reduces Production of Aβ42. **(A)** Schematic representation of *APP* and sequence of *APP* exon 17 and introns 16 and 17, showing the absence of NGG motifs that enable targeting of splicing elements with Sp Cas9. **(B)** Genomic DNA editing rates at the SA of *APP* exon 17 with ABE8e and CBE4max fused with Sp Cas9 or SpRY Cas9 in HEK293T cells; n=3; **, p<0.01; one-tailed t-test. **(C)** SA DNA editing rates accomplished with SpRY Cas9 ABE8e when editing the SA alone or in combination with a panel of sgRNAs targeting the SD. **(D)** SD DNA editing rates with sgRNAs targeting the SD alone or in combination with a sgRNA targeting the SA. Editing at the canonical and cryptic SDs is highlighted. **(E)** *APP* exon 17 exon skipping rates with a sgRNA targeting the SA in combination with one of 3 different sgRNAs targeting the SD measured by NGS. **(F)** qPCR quantification of total *APP* mRNA. **(G,H)** Quantification of *APP* exon 17 exon splicing (top) and schematic representation of the corresponding splicing event (bottom) following targeting the SA with a sgRNA and SD with two different sgRNAs. **(I)** Differential gene expression analysis following targeting of SA and SD of *APP* exon 17 in comparison with control cells. Genes with significant upregulation are represented red (adjusted p<0.05). **(J)** Heatmap with Log_2_ fold expression change compared to control for the most and least differentially expressed genes in DNA damage repair and Alzheimer’s pathways. No genes show statistically differential expression. **(K)** DNA editing at the SA and **(L)** SD of *APP* exon 17 in BE(2)-M17 neuroblastoma cells following enrichment with puromycin. **(M)** exon skipping rates for *APP* exon 17 in puromycin selected pools of BE(2)-M17 neuroblastoma cells. **(N)** Aβ42 levels in BE(2)-M17 cells following skipping of *APP* exon 17 in comparison with control cells using ELISA. n=3; ns, no significance; *, p<0.05; **, p<0.01; ***, p<0.001; One-Way ANOVA, Tukey’s Post Hoc comparing SA/SD to SA and SD for all experiments except RNA-seq. For RNA-seq, differential expression was analyzed with the Limma-voom software package, where expression was measured via a modified t-test with Benjamini-Hochberg adjusted p-values used for p-value cutoffs.

Despite the high rates of editing observed at the SA site for *APP* exon 17, we observed limited skipping (∼0.9%) when targeting the SA alone (**Fig. 4E**). However, when the SD was targeted with each of the 3 different sgRNAs described above, we observed exon skipping (**Fig. 4D-E**). In addition to editing at the SD, we identified two potential cryptic ‘GT’ splice donors that were edited, and their editing appeared to influence exon skipping (**Fig. 4D-E**). While editing at the canonical SDs for sgRNAs SD 1 and SD 2 were similar, sgRNA SD1 edited only the intronic cryptic SD while sgRNA SD2 edited only the exonic cryptic SD. The exon skipping rates were vastly different for sgRNAs SD 1 and 2, 69% and 6.5% respectively, demonstrating that the intronic cryptic SD plays a critical role in *APP* exon 17 splicing. Interestingly, when utilizing dual splice site editing with sgRNAs SA and SD 3 (**Fig. 4D-E**), there was editing at the cryptic intronic SD but not at the canonical SD, yet exon skipping was improved when editing the SA and the intronic ‘GT’ sequence. Furthermore, simultaneously targeting the SA and SD sites improved exon skipping rates for all sgRNA pairs tested, ranging from a slight increase to a synergistic improvement (69% to 75% for SA/SD 1, 6.5% to 24% for SA/SD 2, 15% to 27% for SA/SD 3, p < 0.05 for all) (**Fig. 4E**). Importantly, splice site editing did not significantly (p > 0.05) affect *APP* mRNA expression measured by qPCR, which is critical as *APP* has essential roles in multiple signaling pathways. (**Fig. 4F**). Notably, when characterizing the precision of the splicing events following editing of *APP* exon 17, we observed that simultaneous targeting of both the SA and SD again reduced cryptic splicing events (**Fig. 4G-H**). For example, when the SA was targeted individually, its modification caused a cryptic ‘AG’ motif located between V692 and E693 to be utilized for splicing, leading to a premature termination codon within exon 17 in ∼7% of transcripts, an outcome that was almost completely abolished with dual splice site-editing (SA/SD 1 vs SA, p < 0.0001). When editing the canonical SD with the sgRNA SD 2, a partial intron retention event, whereby the cryptic intronic SD functioned as a canonical SD, was found to occur in 39% of transcripts, further showing the importance of this intronic ‘GT’ site. This aberrant splicing was reduced by over 50% with simultaneous editing of SD and SA (SA/SD 2 vs SD 2, p = 0.0004), which is critical as this cryptic splicing event preserves the protease cleavage sites for the β- and γ-secretases and, therefore, its presence is unlikely to reduce Aβ42. SA/SD 1 achieved the highest overall exon skipping rates and was thus used in subsequent experiments.

To determine if our dual-splice site editing platform influenced overall RNA expression on a transcriptome-wide scale, we performed RNA-seq and differential gene expression (DGE) analysis following targeting of *APP* exon 17.^52^ This analysis revealed differential expression of only 3 genes, *HSPA6*, *ZCCHCH12*, and *RPL21P120* (Benjamini-Hochberg adjusted p < 0.05, **Fig. 4I**). These genes were all upregulated, have no association with DNA damage repair or mismatch repair but have been linked to late-onset AD.^53,54^ No genes were found to be downregulated. Additionally, we analyzed gene expression pathways associated with mismatch repair (KEGG pathway hsa03430) and base excision repair (KEGG pathway hsa03410) and observed no changes in pathway expression for any genes when compared to control cells (DEG defined as a gene with a Benjamini-Hochberg adjusted p > 0.05 for upregulated genes, **Fig. 4 J, S3B,C**), indicating SPLICER did not induce DNA damage repair pathways nor misregulate genes different from the intended target. When analyzing the transcriptomic profile for genes linked with AD (KEGG pathway hsa05010), we found no change in expression associated with pathways implicated in the pathogenesis of AD, including *APP* (DEG defined as a gene with a Benjamini-Hochberg adjusted p > 0.05 for upregulated genes, **Fig. 4I-J, S3D**), further demonstrating that simultaneous SA/SD targeting does not induce misregulation of gene expression.

Next, to evaluate the effect of dual-splice site editing on the formation of Aβ42, we targeted *APP* exon 17 in BE(2)-M17 cells, which are commonly used to model aspects of AD as they produce high levels of the Aβ42 peptide.^55^ Accordingly, we first sought to determine if *APP* could be effectively edited in BE(2)-M17 cells with the lead editing guide candidate identified in the experiments above. Following puromycin enrichment of transfected cells, DNA editing at the SA was ∼60%, while the cryptic intron SD and canonical SD were both edited at ∼100% (**Fig. 4K-L**) and exon skipping when simultaneously targeting the SA and SD sites was ∼90% (**Fig. 4M**). When editing the SA alone, a cryptic ‘AG’ splice acceptor motif was recognized between V692 and E693, like in HEK293T cells, at a rate of 21%. This cryptic splicing was reduced to <1% with simultaneous editing (p < 0.0001, **Fig. S3E**). No other alternative splicing was observed (**Fig. S3E**). We then performed ELISA to quantify expression of Aβ42, measuring a ∼70% decrease in Aβ42 peptide following simultaneous SA/SD editing (SA/SD vs ROSA, p = 0.015) (**Fig. 4N**). Finally, an analysis of editing at potential off-target sites predicted by Cas-OFFinder^56^ revealed no appreciable A>G editing at any computationally predicted sites (p > 0.05, **Fig. S3F**). These results demonstrate that SPLICER can enable *APP* exon 17 skipping and reduce expression of Aβ42.

To provide further validation of the potential of SPLICER, we sought to induce skipping of *APP* exon 17 in a mouse model of AD. We performed these experiments in APP^K670N/M671L^ mice, which harbor a yeast artificial chromosome expressing the full human *APP* gene, including introns, which are not typically included in other mouse strains.^57–59^ To enable delivery by AAV, we used a split-intein BE system that we previously developed^60^ as the size of a full-length BE exceeds the ∼4.7 kb packaging capacity of the AAV vector genome. In this system, the base editor is split into two vectors at Cas9 amino acid residue 712 and the resulting N-terminus and C-terminus Cas9 domains are fused with the N and C *Rhodothermus marinus* inteins, respectively. Upon expression, the inteins dimerize and self-excise, which results in reconstitution of the full length BEs.

We delivered BE-encoding and EGFP-KASH^61^ encoding AAV to four-to seven week-old mice via a stereotaxic injection to the hippocampus, an area of the brain typically affected in AD by the formation of Aβ42 plaques, with AAVrh10, a serotype that has efficiently transduces the hippocampus (**Fig. 5A**).^29,30,62,63^. Expression of BEs and EGFP-KASH were driven by the CAG promoter. At one month after injection, the transduction efficiency measured by fluorescence activated cell sorting (FACS) in harvested hippocampus tissue was ∼23.6% **(Fig. S4A).** In bulk tissue, our deep sequencing analysis revealed DNA editing rates of 6.4% for the SA (**Fig 5B**), 22.4% for the SD, and 23.4% for the intron cryptic SD (**Fig. 5C**), which are consistent with our findings in cultured cells considering our transduction efficiency. Notably, this editing was ∼3-fold higher in FACS sorted nuclei (**Fig. S4B-C**). Importantly, we observed ∼20% skipping of *APP* exon 17 in bulk hippocampus tissue (SA/SD vs ROSA, p = 0.003, unpaired t-test) (**Fig. 5D**). Our deep sequencing analysis further revealed that there was no aberrant cryptic splicing resulting from SA/SD DNA editing, indicating clean, full-length exon 17 skipping achieved with SPLICER editing *in vivo* (**Fig. 5E**).

**Figure 5.**
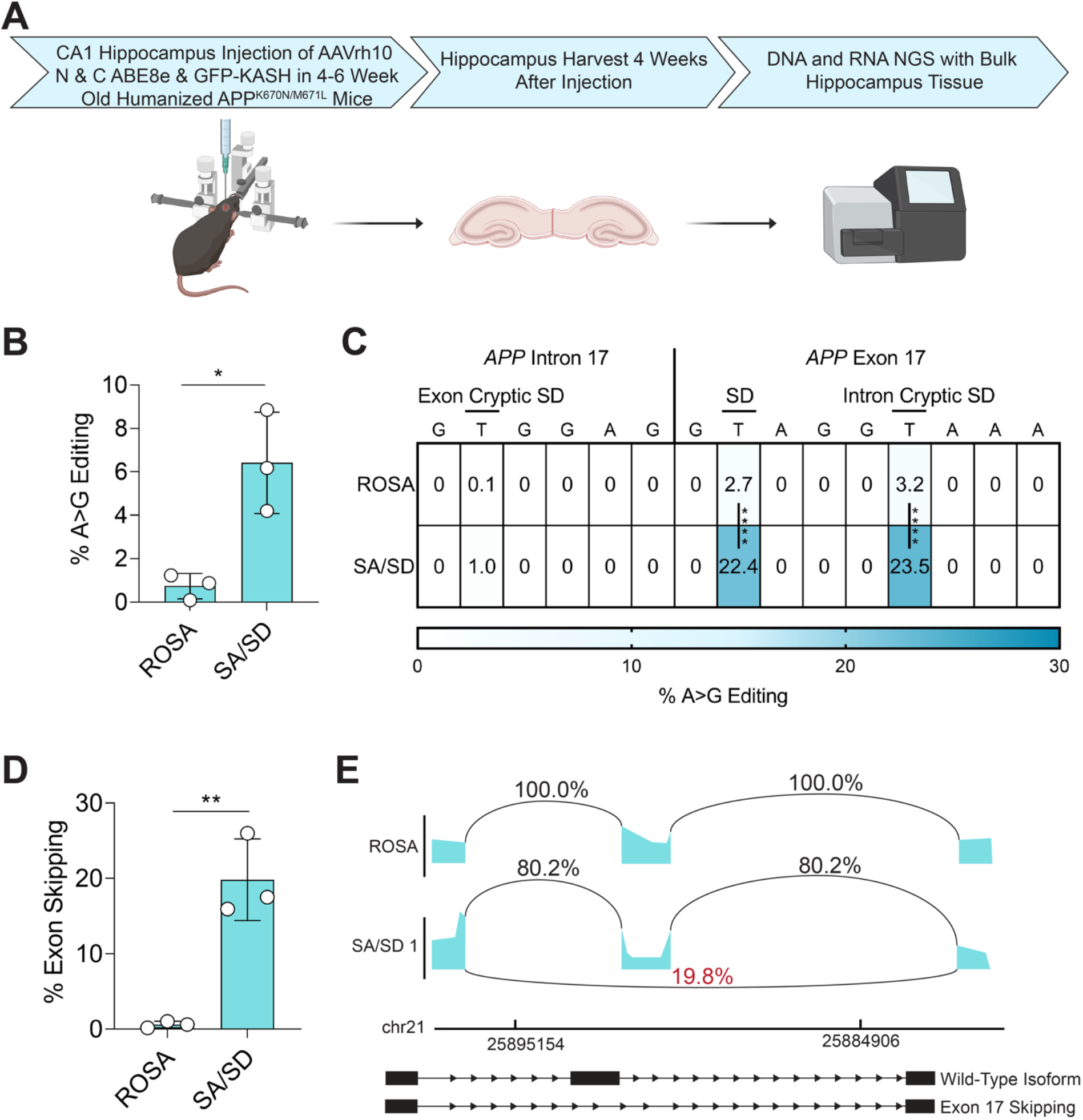
SPLICER Enables Efficient DNA Editing and Full Exon Skipping in A Humanized Mouse Model of Alzheimer’s Disease. **(A)** Experimental workflow of *in vivo* editing following AAVrh10 injection of BEs targeting the SA and SD of *APP* exon 17 in the hippocampus of APP^K670N/M671L^ mice. **(B)** *In vivo* genomic DNA editing of *APP* exon 17 SA and **(C)** SD in bulk hippocampus tissue measured using NGS. **(D)** *In vivo* exon skipping of *APP* exon 17 in bulk hippocampus tissue following treatment with BEs targeting the SA and SD. **(E)** Sashimi plot of splicing events in control mice and mice injected with BEs targeting the SA and SD of *APP* exon 17. n=3 for all groups. *, p<0.05; **, p<0.01; One-way ANOVA, Tukey’s Post Hoc for all experiments.

In conclusion, we developed SPLICER, a method that improves full-length exon skipping by Cas9 base editors. We further demonstrate the therapeutic potential of SPLICER by targeting a high-value target for AD in a mouse model of the neurodegenerative disorder.

## DISCUSSION

In this work we developed SPLICER, a platform for exon skipping with base editors that overcomes the limitations of previously described technologies. This platform utilizes base editors with relaxed PAM requirements fused with different deaminases that, when targeted to both SD and SAs simultaneously, increase exon skipping efficiency and decrease splicing aberrations.

Potential applications of SPLICER include therapeutic exon skipping to restore protein expression^64–68^, disruption of splicing sequences to interrogate and dissect exon splicing, gene knock out by altering the reading frame through exon skipping^15,69^ or massively parallel loss-of-function screens^70^, which also take advantage of gene knock out by shifting the reading frame.

SPLICER provides significant advantages over other methods for exon skipping such as ASOs, which are currently used as gene therapies for treating Duchene muscular dystrophy^71,72^ or retinitis pigmentosa^73,74^, because the transient nature and rapid clearance of ASO requires repeated administration and provides only a temporary benefit with limited activity in the periods of time leading up to the new dosage. The permanent nature of the modifications introduced in genomic DNA by base editors is advantageous, although the transient effects of ASOs are reversible, which could be beneficial if adverse effects occur.

Other gene editing technologies, such as traditional CRISPR-Cas9 nucleases^75,76^ or prime editors,^77,78^ can also be used to disrupt splicing sequences. However, as the knowledge about the potential deleterious effects of DSBs increases, it becomes critical to utilize technologies that minimize DSBs and introduce minimal genotoxicity, which can make base editors preferable over nucleases or prime editors.^1,79,80^

One limitation of base editors is their capacity to introduce bystander mutations, which could hinder therapeutic applications that require precise correction of a single base mutation.^81^ This problem is not critical for exon skipping applications because undesired bystander mutations often occur in intronic sequences where they are unlikely to have a detrimental effect. Further, when they are found in exonic sequences, they are unlikely to have an impact on the protein products, as the exon is skipped.

Guide-dependent and guide-independent off-target effects are important concerns when using base editors.^82,83^ Since SPLICER targets sequences that are conserved in the genome, it is possible that the sgRNAs utilized may be at a higher risk for off-target effects. However, given that only 2 base pairs within SD or SAs are highly conserved, and the guides themselves almost invariably target partly exonic sequences, which are less conserved, we unsurprisingly did not observe any off-target mutations at computationally predicted off-target sites for the sgRNAs targeting *APP* exon 17. Nonetheless, the analysis of off-target effects was limited to only for five predicted off-target sites and further analysis using unbiased methods for detecting off-target mutations such as EndoV-seq^84^ will be needed to get a more comprehensive view of off-target editing for dual splice site editing for *APP* exon 17. Guide-independent off-target effects are derived from the DNA binding capability of the deaminases and occur more frequently when using CBE than ABEs.^83,85,86^ While newer generations of highly specific CBEs have been developed^40–42^, we generally utilize ABEs whenever possible, which is advantageous not only because of their specificity but also because of their higher editing activity. Importantly, a CBE was recently developed from ABEs^42^, which has improved specificity and has proven to be effective for inducing exon skipping in our work.

SPLICER was not sufficient to accomplish skipping of all targeted exons, including AHCY exon 9 (**Fig. 1**). One possibility for this lack of improved exon skipping is the presence of exon splice enhancer (ESE) sequences, which bind to Ser/Arg rich (SR) proteins that recruit spliceosome proteins.^87,88^ We studied the presence of ESE utilizing ESEfinder^88^ and determined that, within AHCY exon 9, there are motifs predicted to function as splice enhancers (high affinity to SF2/ASF: score 3.76, SC35: score 3.69, SC35: score 4.14, SRp40: score 3.88, SRP55, score 4.69)^88^, which may explain why this particular exon appeared resistant to exon skipping. Interestingly, Qiu and coworkers have shown recently that ESE can be targeted with BEs to accomplish exons skipping.^89^ While their results are intriguing, the approach has high potential to introduce exonic mutations without accomplishing exon skipping, which could lead to adverse effects.

One important caveat of our analysis is that we performed targeted PCR in combination with NGS to study skipping outcomes. However, in instances in which the outcome is a frame shift followed by nonsense mediated decay it is likely that the mRNA is degraded too quickly to be detected using our approach. Similarly, in cases of intron retention where the amplicon is too large to be detected by the chosen PCR primers, it is likely that our approach failed, resulting in underestimation of this splicing outcome. Future studies utilizing long-read analysis will be needed to further improve our understanding of exon skipping.

Lastly, we show the application of SPLICER for skipping *APP* exon 17, an approach with potential therapeutic value in Alzheimer’s disease. By skipping exon 17, given that the binding site for the secretases is lost, generation of Aβ42 decreases. Previous attempts to skip this exon in our lab had been unsuccessful despite high editing rates at the SA, as a lack of tools for targeting additional splice sites prevented further optimization of the approach. Using SPLICER, we have identified multiple sgRNAs with different editing profiles that have contributed to our understanding of the splicing events governing this transcript. We observed that editing the canonical SD activates a cryptic SD that prevents exon skipping and only when simultaneously editing both the canonical SD and cryptic SD sites with SPLICER we were able to effectively induce exon skipping. Additionally, *APP* exon 17 skipping was further enhanced by the simultaneous disruption of the SA, an event that reduced Aβ42 production *in vitro*. SPLICER had no effect on *APP* mRNA levels of expression, as determined by qPCR and RNA-seq, however, RNA-seq did reveal that *APP* exon 17 disruption affected the expression of the genes *HSPA6*, *ZCCHCH12*, and *RPL21P120*. Interestingly, the expression of these genes has been shown to be modulated in late-onset forms of AD, indicating that the Aβ42 may play a role in regulating expression of these genes, further illustrating the complex role of Aβ42^53,54^. Critically, SPLICER functions efficiently *in vivo* for targeting *APP* exon 17, accomplishing 20% exon skipping in a humanized *APP* mouse model when delivered by AAVrh10. While the skipping rate *in vivo* was approximately three-fold lower than the observed *in vitro* efficiency, when taking into consideration the apparent transduction efficiency, which we measured as ∼23.6%, this result emphasizes the potential of the approach, which we anticipate could be further improved following optimization of delivery.

In summary, we report the development of the SPLICER toolbox, which enables targeting of essentially any exon and improves skipping purity outcomes via simultaneous editing of multiple splicing elements. We demonstrate the *in vivo* applicability of SPLICER in a humanized mouse model of Alzheimer’s disease and anticipate that this technique will enable a range of broad applications.

## METHODS

### Plasmids and Cloning

The plasmid encoding the U6-sgRNA expression cassette was obtained from Addgene (#47108). The full-length SpRY CBE4max construct was purchased from Addgene (Plasmid #139999). The full-length SpRY ABE8e plasmid was generated through Gibson Assembly of gBlock Gene Fragments (Integrated DNA Technologies) containing the adenosine deaminase in Addgene Plasmid #138489 and the SpRY CBE4max backbone. To build the ABE and CBE constructs consisting of different deaminases described in **Fig. 1**, first SpRY mutations were cloned via Gibson Assembly of gBlock Gene Fragments (Integrated DNA Technologies) into BE-expressing plasmids described in our previous work.^18,90^. Then, different deaminases were cloned into the plasmids via Gibson Assembly of gBlock Gene Fragments. For experiments in BE(2)-M17 cells, which utilized a plasmid encoding SpRYABE8e-T2A-Puro, SpRY mutations, ABE8e deaminase, and the puromycin resistance gene were cloned via Gibson Assembly of gBlock Gene Fragments into the lentiviral expression plasmid pLenti-EFS-FNLS-P2A-BlastR.^11^ Plasmids encoding the split AAV SpRY ABE8e constructs were cloned via Gibson Assembly of gBlock Gene Fragments into our previously described AAV split Cas9 BE3 plasmids to insert SpRY mutations as well as to replace APOBEC1 with ABE8e.^90^ All amino acid sequences from these plasmid constructs are provided in **Table S1**.

All oligonucleotides used in this work were obtained from Integrated DNA Technologies. The oligonucleotides for sgRNA generation were hybridized, phosphorylated and cloned following the U6 sgRNA vector using BbsI, Bsa1, of BsmBI sites with sequences are provided in **Table S2**.

### Cell Culture and Transfections

The cell line HEK293T was obtained from the American Type Culture Collection (ATCC) and was maintained in DMEM supplemented with 10% fetal bovine serum and 1% penicillin/streptomycin at 37°C with 5% CO2. BE(2)-M17 cells was obtained from the ATCC and maintained in 1:1 EMEM:Ham’s F12K supplemented with 10% fetal bovine serum and 1% penicillin/streptomycin at 37°C with 5% CO2.

HEK293T cells were transfected in 24-well plates with Lipofectamine 2000 (Invitrogen) via reverse transfection following the manufacturer’s instructions. The amount of DNA used for lipofection was 1 μg per transfection. In experiments with split base editors, 500 ng of each split were used. For simultaneous targeting experiments in HEK293Ts, 500 ng of plasmid encoding full-length BEs were transfected with 500 ng of plasmid with the sgRNA expression cassette when one sgRNA was expressed. Whenever two separate sgRNAs were transfected, 333.33 ng of plasmid encoding BEs was used with 333.33 ng of each sgRNA cassette. Transfection efficiency was routinely higher than 90% for HEK293T cells as determined by fluorescent microscopy following delivery of 1 µg of a GFP expression plasmid.

### AAV Vector Production

AAV was produced according to the protocol described previously.^91^ HEK293T cells were seeded onto 15-cm plates to be 80% confluent after 16 hours and maintained in DMEM supplemented with 10% (v/v) FBS and 1% (v/v) penicillin/streptomycin. After 16 hours, cells were transfected with 65 μg total of AAV vector plasmid (pAAV-CAG-N-ABE8e-SpRY-APPex17sgRNA, or pAAV-CAG-C-ABE8e-SpRY-APPex17sgRNA, or pAAV-CAG-GFP-KASH), pAAVrh10, and pHelper in a 1:1:1 mol ratio using PEI-max (pH=8). XmaI digestion was used to confirm the integrity of the pAAV plasmids before transfection. Cell media was replaced 4 hours after transfection. Cells were harvested 72 hours after transfection by manual dissociation using a cell scraper and centrifuged at 1500 x g for 5 min at room temperature. The supernatant was collected into a fresh tube and mixed with 40% polyethylene glycol 8000 (Thermo Fisher) solution in a ratio of 4:1 by volume and stored overnight at 4°C. Cell pellets were then resuspended in 2 mL of lysis buffer (50 mM Tris-HCl and 150 mM NaCl, pH 8.0) per plate. The next day, the supernatant solution was centrifuged at 3,000 RPM for 30 minutes at 4°C. The pellets of the supernatant were resuspended in the same lysis buffer as described above and mixed with their respective cell lysates. These cell suspensions were then frozen in liquid Nitrogen and thawed at 37°C, for three freeze-thaw cycles to extract AAV vectors. Supernatants were then treated with 0.5% Triton X-100 (Thermo Fisher) and 50 units/mL Benzonase (Merck) shaking at 37°C for 1 hour. We then centrifuged the lysate at 10,000 x g for 15 minutes at room temperature. The resulting supernatant was overlaid onto an iodixanol density gradient using 15%, 25%, 40% and 60% Opti-Prep solution (Sigma-Aldrich) and the virus was isolated by ultracentrifugation at 58,400 RPM at 18°C. This step was repeated to perform a second gradient purification using iodixanol fractions of 30%, 40% and 60%. Following extraction, AAV was filter-dialyzed with 1X PBS containing 0.001% Tween-20 using an Amicon Ultra 100 kDa MWCO column (Merck). The titer of the purified AAV was determined, post-treatment with DNase I (Millipore Sigma), by quantitative real-time PCR using primers that amplify the bGH PolyA sequence (**Table S3**) and the SsoFast Evagreen mix (Bio-Rad). The virus was stored at -80°C.

### RT-PCR

RNA was harvested from cell pellets using the RNeasy Plus Mini Kit (Qiagen) according to manufacturer’s instructions. cDNA synthesis was performed using the qScript cDNA Synthesis Kit (Quanta Biosciences) from 1 µg of RNA with cycling conditions performed as directed by the supplier. PCR was performed using KAPA2G Robust PCR kits from Kapa Biosystems. The 25 µL reactions used 25 ng of cDNA, Buffer A (5 µL), Enhancer (5 µL), dNTPs (0.5 µL), 10 µM forward primer (1.25 µL), 10 µM reverse primer (1.25 µL), KAPA2G Robust DNA Polymerase (0.5 U) and water (up to 25 µL). Cycling parameters were used a recommended by the manufacturer. The PCR products were visualized in 2% agarose gels stained with ethidium bromide and images were captured using a ChemiDoc-It2 (UVP). The DNA sequences of the primers for each target are provided in **Table S3**.

### Densitometry Analysis

Skipping efficiencies for screening of simultaneous sgRNA candidates were determined by densitometry analysis of the PCR products obtained from RT-PCR and analyzed by agarose gel electrophoresis using ImageJ software. After subtracting background noise, band intensity was compared using the following formula: % exon skipping = (Skipped Band Intensity)/(Non Skipped Band Intensity + Skipped Band Intensity) where band intensity is the sum of each pixel grayscale value within the selected area of the band.

### Analysis of DNA Editing Rates

Genomic DNA was isolated using a DNeasy Blood and Tissue Kit (Qiagen) and PCR amplification was performed with KAPA2G Robust PCR kits (KAPA Biosystems) as described above, using 20-100 ng of template DNA and primers listed in **Table S3**.

Sanger sequencing of the PCR amplicons was performed by the Roy J. Carver Biotechnology Centerat the University of Illinois at Urbana-Champaign. Base editing efficiencies were estimated by analyzing sequencing traces using EditR with the primers listed in **Table S3**.

### Next-Generation Sequencing for RNA Exon Skipping and DNA Amplicons

After cDNA synthesis (qScript, QuantaBio), amplicons were generated using KAPA HiFi HotStart (Roche), according to manufacturer’s instructions with primers containing overhangs compatible with Nextera XT indexing (IDT). Following validation of the quality of PCR products by gel electrophoresis, the PCR products were isolated using an AMPure XP PCR purification beads (Beckman Coulter). Indexed amplicons were then generated with a Nextera XT DNA Library Prep Kit (Illumina) and pooled. Libraries were sequenced with a MiSeq Nano Flow Cell for 251 cycles from each end of the fragment using a MiSeq Reagent Kit v2 (500-cycles). FASTQ files were created and demultiplexed using bcl2fastq v2.17.1.14 Conversion Software (Illumina). Deep sequencing was performed by the W.M. Keck Center for Comparative and Functional Genomics at the University of Illinois, Urbana, IL.

Exon skipping rates were quantified using the STAR RNA-Seq aligner on Galaxy. Forward and reverse reads were combined and aligned to the human reference genome (GRCh38) using STAR. 2-pass mapping was used for splice junction analysis with a MAPQ value of 60 for .bam files. Splice junctions formed were determined from STAR SJ.out.tab files where the percentage of a junction event was defined as the number of reads for the target junction divided by the total number of events at the specified junction. Sashimi plots for splice junctions were generated from the Integrative Genomics Viewer (IGV’s) sashimi plot tool using the .bam and .bai files outputted from RNA-STAR. These sashimi plots were illustrated in this manuscript by tracing the images generated from IGV in Adobe Illustrator.

For DNA amplicons, samples were isolated with the DNeasy Blood and Tissue Kit and the RNeasy Plus Mini Kit per manufacturer’s specifications. Amplicons were generated by PCR with KAPA HiFi Hotstart using primers containing overhangs compatible with Nextera XT indexing primers listed in **Table S3**. Amplification, indexing, pooling, and sequencing was performed as described above. Base editing rates were quantified using CRISPResso2. Reads with average phred scores below 30 were removed, and remaining reads were aligned to the expected amplicon sequences.

### RNA-Seq

RNA was harvested as previously mentioned in this manuscript. Following analysis of RNA on a 1% agarose gel for RNA purity, RNA seq libraries were prepped with the TrueSeq® mRNA library prep kit (Illumina) and samples were sequenced with a NovaSeq 6000 system (Illumina). For analysis, a quality check performed using FASTQC showed that average per-base read quality scores in all samples were above 34 and no adapter sequences were found, indicating high quality reads, requiring no trimming. The reads were then mapped to the human genome (hg38) using HISAT2, with over 96% of reads being mapped uniquely. These mapped reads were counted across human genes with featureCounts, using annotations of hg38 obtained from GENCODE. About 60% of reads were assigned to genes. Differential expression analysis was performed using limma-voom which first calculates TMM normalized counts-per-million (CPM) to account for compositional bias. In addition, genes expressed at very low levels (CPM lower than 0.5) were filtered out before performing differential expression analysis. For DEG, a DEG was defined as a gene with a Benjamini-Hochberg adjusted p > 0.05.

### RT-qPCR

RNA harvesting and cDNA synthesis was performed as described previously in this manuscript. For qPCR, 50 ng of cDNA was used per sample. qPCR was performed with SsoFast™ EvaGreen® Supermix (Biorad) following manufacturer recommendations for both reaction setup and cycling conditions. 500 nM of qPCR primers were used per sample, and GAPDH was used as the housekeeping gene. Thermocycling and cycle threshold (ct) measurements was conducted on the CFX96™ qPCR instrument (Biorad). Gene expression levels were analyzed from ct values of the target genes and relative expression was quantified using the double-delta ct method.^92^ All primers used for qPCR are listed under **Table S3**.

### ELISA

To generate cells with editing at *APP* exon 17, BE(2)-M17 cells were transfected via reverse transfection with lipofectamine 2000 containing 1 ug of pEFS-SpRY-ABE8e-T2A-Puro, a plasmid which also contained a U6 driven sgRNA expression cassette. For samples expressing multiple sgRNAs, 500 ng of each plasmid was used. 24 hours after transfection, cell media was exchanged with media supplemented with 1 µg/mL puromycin and incubated for 36 hours.

For ELISA, cells were plated onto 10 cm dishes and grown to 100% confluency. After cells reached confluency, cell media was replaced and incubated for an additional 24 hours. Media was then replaced with 10 mL OPTI-MEM supplemented with 1% penicillin/streptomycin and cells were incubated for 48 hours. Cell supernatant was collected into 1.5 mL tubes, supplemented with 1X Halt™ Protease Inhibitor Cocktail (Thermo™ Scientific 78430) and centrifuged at 1400 rpm for 1 minute. Supernatant was collected, and an Aβ42 ELISA was performed on undiluted samples using the Ultrasensitive Aβ42 Human ELISA Kit (Thermo™ Scientific KHB3544) according to the manufacturer’s instructions. Standards were diluted in OPTI-MEM supplemented with 1% penicillin/streptomycin and 1X Halt™ cocktail.

For total protein analysis, adhered cells were lysed with 1X RIPA buffer on ice with 1X Halt™ cocktail and incubated on ice for ten minutes. Cells were spun at 10,000 rpm for 10 minutes, and total protein content in the supernatant was quantified with the Pierce™ Bradford Protein Assay Kit (Thermo™ Scientific 23200) following manufacturer instructions.

After samples were quantified via ELISA, the concentration of Aβ42 was normalized to the total protein content of each sample by dividing each Aβ42 concentration by the total protein concentration. The value generated by each sample was normalized to its respective ROSA positive control.

### Stereotaxic injections

All animal procedures were approved by the Illinois Institutional Animal Care and Use Committee at the University of Illinois and conducted in accordance with the National Institutes of Health (NIH) Guide for the Care and Use of Laboratory Animals. Four to six week old transgenic B6.129-Tg(APPsw)40Btla/Mmjax mice (Jackson Laboratory #0034831-JAX) were injected with 5 x 10^9^ vector genomes of each of AAVrh10-pAAV-CAG-N-ABE8e, pAAVrh10-CAG-C-ABE8e, and 5 x 10^8^ vector genomes of AAVrh10-pAAV-CAG-KASH in 2 μl of PBS with 0.001% Tween-20 into each half of the hippocampus at coordinates -1.8 AP, ±1.5 ML, and 0.45 DV.

### Tissue Harvesting and RNA Purification

Mice were anesthetized using 3% isoflurane delivered through vaporizer in a closed chamber and transcardially perfused using 1 X PBS. The hippocampus was dissected, divided in 2 halves and each half was stored via flash freezing or in RNAlater (Invitrogen, AM7026).

For FACS, nuclei isolation was performed as previously described.^93^ Briefly, tissues were homogenized in NF1 buffer using the KIMBLE Dounce Tissue Grinder (Sigma-Aldrich) per the manufacturer’s instructions, strained through a 70 µm strainer into a 50 mL conical tube underlayed with a 1.2 M sucrose cushion, and centrifuged at 3,900G for 30 minutes at 4°C. The supernatant was discarded and the pelleted nuclei were resuspended in 10 mL of NF1 buffer. Samples were spun 2X at 1,600G for 5 minutes at 4°C and resuspended in 2 mL of FANS buffer. Cells were strained through a 35 µm strainer and incubated at 37°C for 30 minutes with 0.5 µL Vybrant™ Dyecycle™ Ruby strain (Invitrogen, V10309) per mL of FANS buffer.

Harvested nuclei were sorted using a ThermoFisher Bigfoot Spectral Cell Sorter Cell Sorter (Roy J. Carver Biotechnology Center, University of Illinois, Urbana, IL). Cells were collected in FANS buffer. At least 10,000 cells or nuclei were sorted for each sample. Collected nuclei were diluted in FANS buffer with 1% wt/vol (g/mL) BSA and spun at 1,600G for 15 minutes at 4°C. Supernatant was aspirated, and pelleted nuclei were isolated via the DNeasy Blood and Tissue Kit To perform downstream analysis on whole hippocampus samples, DNA and RNA samples were isolated with the DNeasy Blood and Tissue Kit and the RNeasy Plus Mini Kit per manufacturer’s specifications, respectively. Prior to purification, brain tissue samples were homogenized in 1X PBS using a KIMBLE Dounce Tissue Grinder, diluted in the first buffer of each respective kit, and further isolated following the standard kit protocols.

For NGS on all DNA samples, Amplicons were generated by PCR with KAPA HiFi Hotstart using primers containing overhangs compatible with Nextera XT indexing primers listed in **Table S3**. Amplification, indexing, pooling, and analysis of base editing rates was performed as described in the “*Next-Generation Sequencing for RNA Exon Skipping and DNA Amplicons*” section. For RNA analysis of exon skipping, cDNA synthesis and amplification for NGS was performed as previously described in the manuscript, however, libraries or cDNA were sequenced with the MiSeq Bulk flow cell for 275 cycles from each end of the fragment using a MiSeq Reagent Kit v3 (500-cycles). Exon skipping was quantified with RNA-STAR as described above.

### Statistical analysis

GraphPad Prism version 9.1 (GraphPad Software, Inc.) software was used for statistical analysis. All experiments consisted of independent replicates. Test groups were compared using either student’s t-test or a One-Way ANOVA with Tukey’s post-hoc analysis. Statistical methods used for RNAseq are described within the section titled “*RNA-Seq”*.

## DATA AVAILABILITY STATEMENT

All data necessary to evaluate the conclusions is presented within the manuscript or in supplementary material. Deep sequencing files have been deposited in the Gene Expression Omnibus, accession number 246588.

## ACKNOWLEDGMENTS

We thank the DNA Services staff of the Roy J. Carver Biotechnology Center at the University of Illinois, particularly Alvaro Hernandez and Chris Wright, for their support with DNA and RNA sequencing. We thank Siva Mayandi of the Roy J. Carver Biotechnology Center Cytometry and Microscopy to Omics facility at the University of Illinois for his support with FACS. This work was supported by the National Institutes of Health (1U01NS122102, 1R01NS123556, 1R01GM141296, 1R01GM127497, 1R01GM131272), the Muscular Dystrophy Association (MDA602798), the American Heart Association (17SDG33650087), the Parkinson’s Disease Foundation (PF-IMP-1950) and the Simons Foundation (887187). J.W was supported by the Northwestern University Clinical and Translational Science Institute, grant UL1TR001422. A.M was supported by the National Institute of Biomedical Imaging and Bioengineering of the National Institutes of Health under Award Number T32EB019944. The content is solely the responsibility of the authors and does not necessarily represent the official views of the National Institutes of Health.

## AUTHOR CONTRIBUTIONS

A.M., S.S., T.G. and P.P. conceived of the study; A.M., S.S., J.W., G.E., M. G., D.S., W.S.W, N.G., M.G., M.S., A.Z, and H.M. designed and performed experiments; A.N. and S.M. assisted with bioinformatic analysis of RNAseq data. A.M., T.G and P.P. wrote the manuscript with input from all authors.

