## Supplemental data for "SPLICER: A Highly Efficient Base Editing Toolbox That Enables *In Vivo* Therapeutic Exon Skipping"

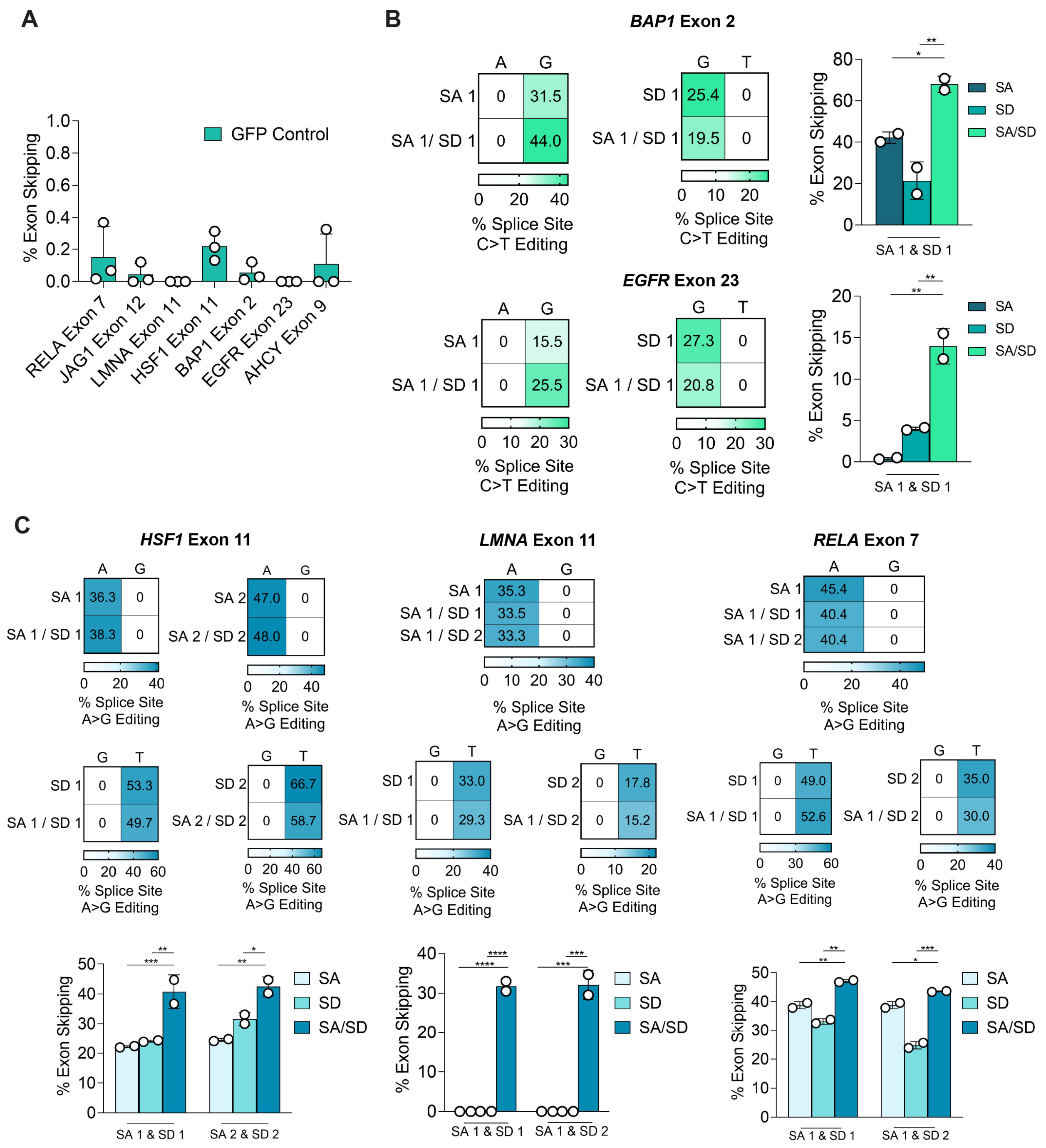


**Figure S1. SPLICER Enhances Exon Skipping. (A)** Quantification of exon skipping by NGS in samples transfected with a non-targeted plasmid control samples transfected controls at targeted exons; n=3. **(B)** DNA editing and RNA exon skipping following targeting of *BAP1* exon 2 (top panel) and *EGFR* exon 23 (bottom panel) with CBEs. **(C)** DNA editing and RNA exon skipping following targeting of *RELA* exon 7, *LMNA* exon 11, *HSF1* exon 11 with ABEs. Exon skipping rates were measured by RT-PCR gel densitometry. n=2; *, p<0.05; **, p<0.01; ***, p,0.001; ****, p<0.0001; One-Way ANOVA, Tukey's Post Hoc comparing SA/SD to SA and SD.

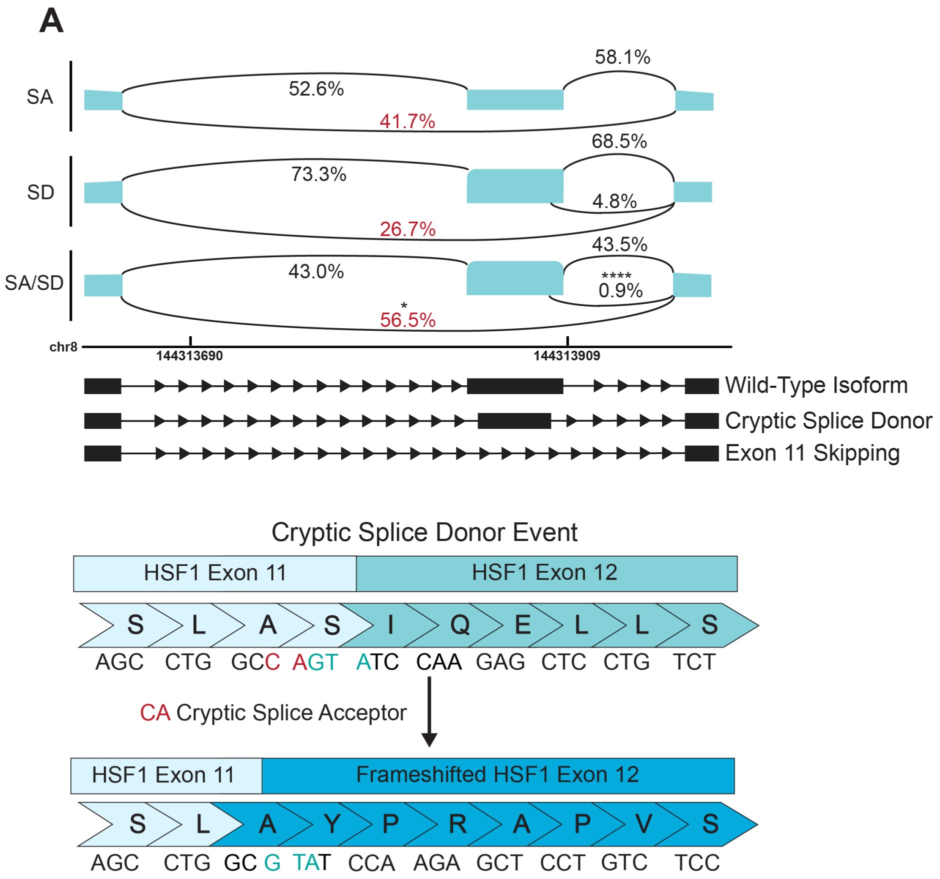


**Figure S2. Reduction of Cryptic Splicing for *HSF1* Exon 11 with SPLICER.** Sashimi plot showing a cryptic splice donor event following target the SA or SD individually or simultaneously (top) leading to a frameshifted protein starting at exon 12. *, p<0.05; ****, p<0.0001; One-Way ANOVA, Tukey's Post Hoc comparing SA/SD to SA and SD.

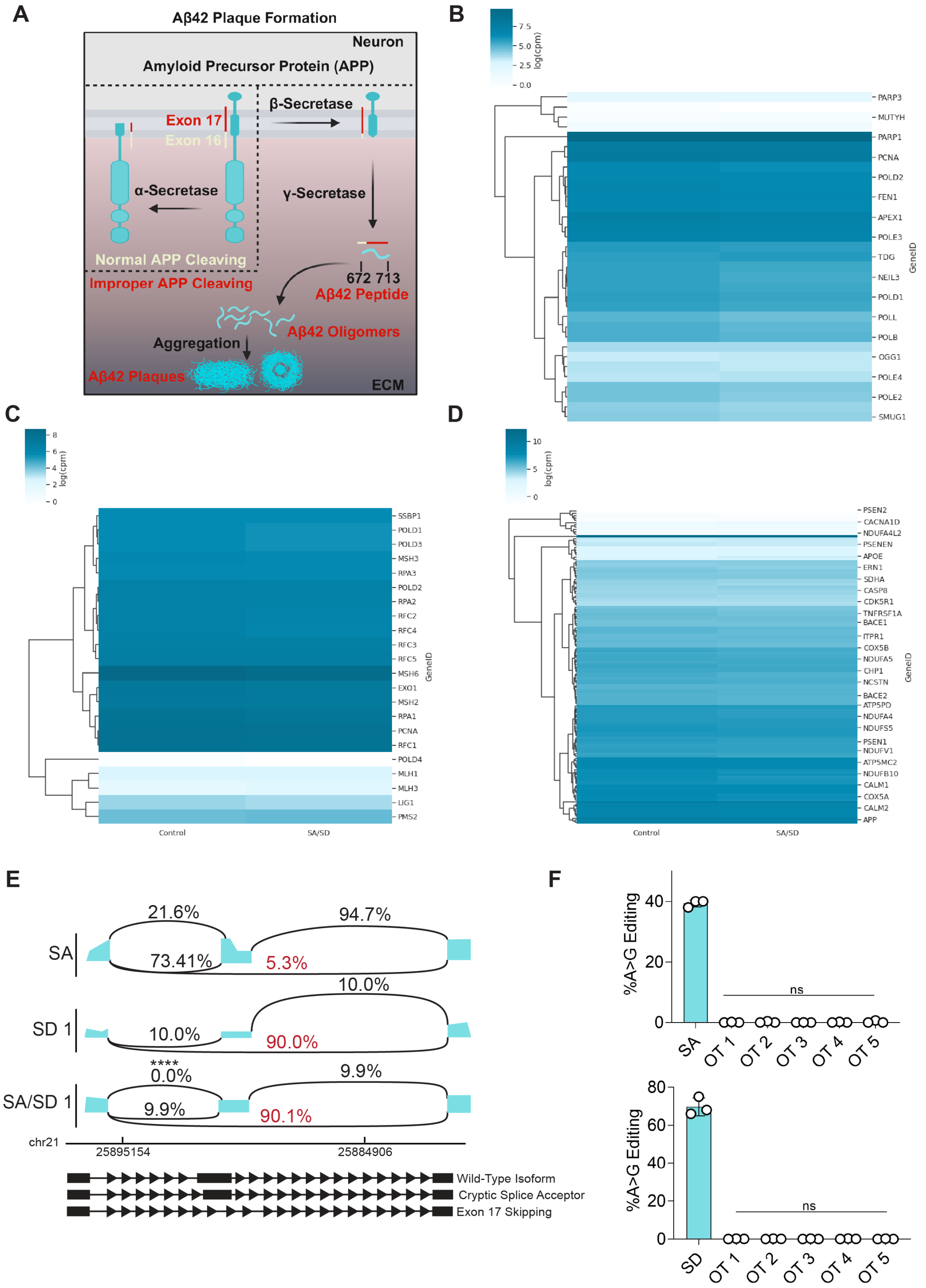


**Figure S3. Analysis of Global Changes in Patterns of mRNA Expression and Off-target Editing Following Induction of Exon Skipping with SPLICER. (A)** Schematic illustrating the pathogenic events leading to formation of plaques in Alzheimer's disease through secretase-mediated cleavage of the APP protein at exon 17. **(B-D)** counts-per-million (cpm) heatmap from RNA-seq data showing no change of the expression levels for genes within the **(B)** base excision repair pathway (KEGG pathway hsa03430), (**C**) mismatch repair pathway (KEGG pathway hsa03410), and **(D)** Alzheimer's disease pathway (KEGG pathway hsa05010) in cells treated with BEs in comparison with control cells. **(E)** *APP* exon 17 exon skipping rates in puromycin-selected BE(2)-M17 cells following treatment with BEs targeting the SA, the SD or both. (F) Analysis of off-target effects of sgRNAs targeting the SA (top panel) or SD (bottom panel) of *APP* exon 17 at 5 computationally predicted sites using NGS. n=3; ns, no significance; ****, p<0.0001; One-Way ANOVA, Tukey's Post Hoc comparing SA/SD to SA and SD.

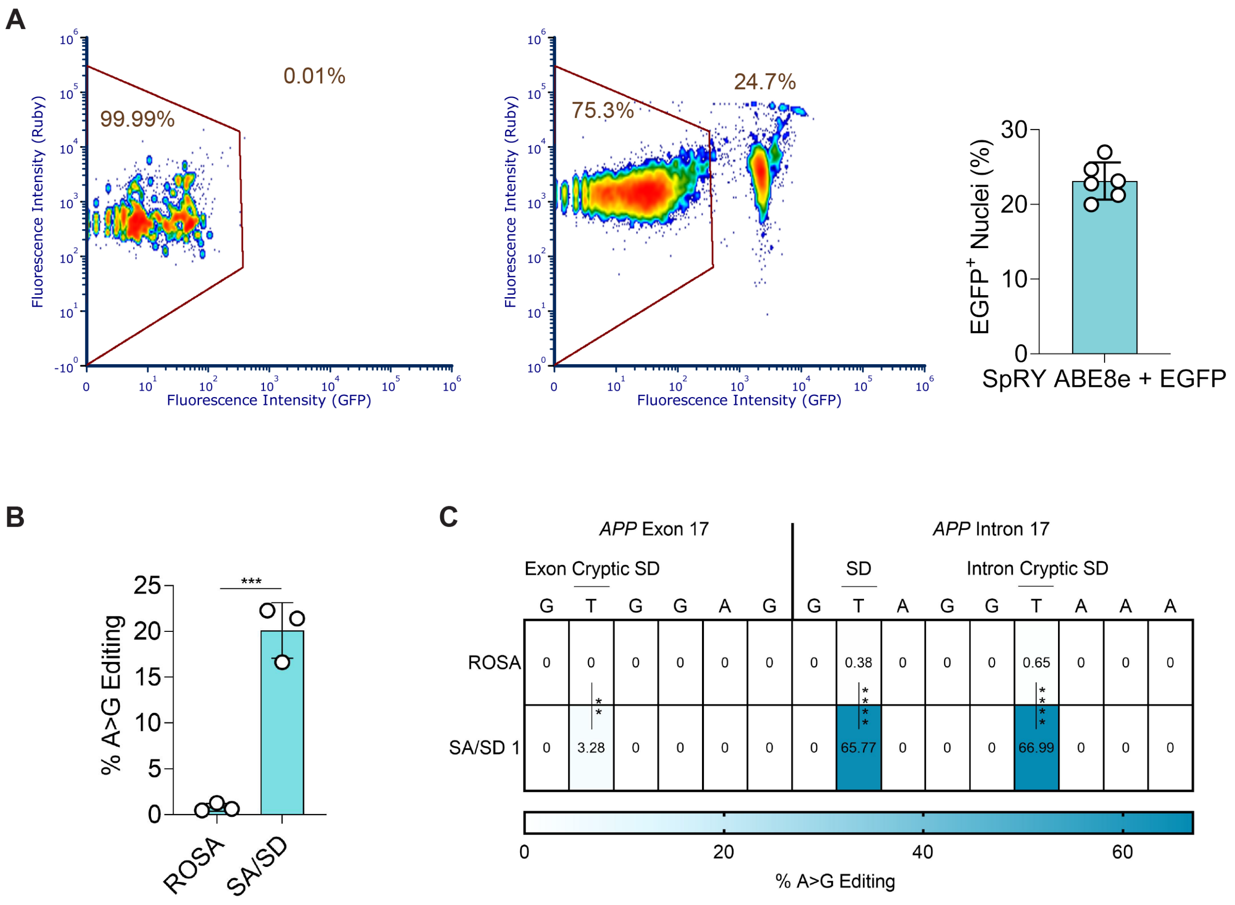


**Figure S4. Transduction Efficiency and DNA Editing in the Hippocampus Following Nuclei Enrichment via FACS. (A)** AAV transduction efficiency in the hippocampus isolated from mice treated with EGFP-KASH BE targeted to the ROSA26 locus or *APP* exon 17 following FACS. **(B)** DNA editing rates at the SA and SD in FACS-sorted nuclei. n=3; **, p<0.01; ***, p,0.001; ****, p<0.0001; One-Way ANOVA, Tukey's Post Hoc.

**Table S1**. Amino Acid Sequence of Cas9 Base Editors

Full Length SpRY ABE8e

MKRTADGSEFESPKKKRKVSEVEFSHEYWMRHALTLAKRARDEREVPVGAVLVLNNRVIGEG
WNRAIGLHDPTAHAEIMALRQGGLVMQNYRLIDATLYVTFEPCVMCAGAMIHSRIGRVVFGVR
NSKRGAAGSLMNVLNYPGMNHRVEITEGILADECAALLCDFYRMPRQVFNAQKKAQSSINSGG
SSGGSSGSETPGTSESATPESSGGSSGGSDKKYSIGLAIGTNSVGWAVITDEYKVPSKKFKVL
GNTDRHSIKKNLIGALLFDSGETAEATRLKRTARRRYTRRKNRICYLQEIFSNEMAKVDDSFFHR
LEESFLVEEDKKHERHPIFGNIVDEVAYHEKYPTIYHLRKKLVDSTDKADLRLIYLALAHMIKFRG
HFLIEGDLNPDNSDVDKLFIQLVQTYNQLFEENPINASGVDAKAILSARLSKSRRLENLIAQLPGE
KKNGLFGNLIALSLGLTPNFKSNFDLAEDAKLQLSKDTYDDDLDNLLAQIGDQYADLFLAAKNLS
DAILLSDILRVNTEITKAPLSASMIKRYDEHHQDLTLLKALVRQQLPEKYKEIFFDQSKNGYAGYI
DGGASQEEFYKFIKPILEKMDGTEELLVKLNREDLLRKQRTFDNGSIPHQIHLGELHAILRRQED
FYPFLKDNREKIEKILTFRIPYYVGPLARGNSRFAWMTRKSEETITPWNFEEVVDKGASAQSFIE
RMTNFDKNLPNEKVLPKHSLLYEYFTVYNELTKVKYVTEGMRKPAFLSGEQKKAIVDLLFKTNR
KVTVKQLKEDYFKKIECFDSVEISGVEDRFNASLGTYHDLLKIIKDKDFLDNEENEDILEDIVLTLT
LFEDREMIEERLKTYAHLFDDKVMKQLKRRRYTGWGRLSRKLINGIRDKQSGKTILDFLKSDGF
ANRNFMQLIHDDSLTFKEDIQKAQVSGQGDSLHEHIANLAGSPAIKKGILQTVKVVDELVKVMG
RHKPENIVIEMARENQTTQKGQKNSRERMKRIEEGIKELGSQILKEHPVENTQLQNEKLYLYYL
QNGRDMYVDQELDINRLSDYDVDHIVPQSFLKDDSIDNKVLTRSDKNRGKSDNVPSEEVVKKM
KNYWRQLLNAKLITQRKFDNLTKAERGGLSELDKAGFIKRQLVETRQITKHVAQILDSRMNTKY
DENDKLIREVKVITLKSKLVSDFRKDFQFYKVREINNYHHAHDAYLNAVVGTALIKKYPKLESEFV
YGDYKVYDVRKMIAKSEQEIGKATAKYFFYSNIMNFFKTEITLANGEIRKRPLIETNGETGEIVWD
KGRDFATVRKVLSMPQVNIVKKTEVQTGGFSKESILPKRNSDKLIARKKDWDPKKYGGFLWPT
VAYSVLVVAKVEKGKSKKLKSVKELLGITIMERSSFEKNPIDFLEAKGYKEVKKDLIIKLPKYSLFE
LENGRKRMLASAKQLQKGNELALPSKYVNFLYLASHYEKLKGSPEDNEQKQLFVEQHKHYLDE
IIEQISEFSKRVILADANLDKVLSAYNKHRDKPIREQAENIIHLFTLTRLGAPRAFKYFDTTIDPKQY
RSTKEVLDATLIHQSITGLYETRIDLSQLGGDSGGSKRTADGSEFEPKKKRKV*

Full Length SpRY CB4max

MKRTADGSEFESPKKKRKVSSETGPVAVDPTLRRRIEPHEFEVFFDPRELRKETCLLYEINWG
GRHSIWRHTSQNTNKHVEVNFIEKFTTERYFCPNTRCSITWFLSWSPCGECSRAITEFLSRYPH
VTLFIYIARLYHHADPRNRQGLRDLISSGVTIQIMTEQESGYCWRNFVNYSPSNEAHWPRYPHL
WVRLYVLELYCIILGLPPCLNILRRKQPQLTFFTIALQSCHYQRLPPHILWATGLKSGGSSGGSS
GSETPGTSESATPESSGGSSGGSDKKYSIGLAIGTNSVGWAVITDEYKVPSKKFKVLGNTDRH
SIKKNLIGALLFDSGETAERTRLKRTARRRYTRRKNRICYLQEIFSNEMAKVDDSFFHRLEESFL
VEEDKKHERHPIFGNIVDEVAYHEKYPTIYHLRKKLVDSTDKADLRLIYLALAHMIKFRGHFLIEG
DLNPDNSDVDKLFIQLVQTYNQLFEENPINASGVDAKAILSARLSKSRRLENLIAQLPGEKKNGL
FGNLIALSLGLTPNFKSNFDLAEDAKLQLSKDTYDDDLDNLLAQIGDQYADLFLAAKNLSDAILLS
DILRVNTEITKAPLSASMIKRYDEHHQDLTLLKALVRQQLPEKYKEIFFDQSKNGYAGYIDGGAS
QEEFYKFIKPILEKMDGTEELLVKLNREDLLRKQRTFDNGSIPHQIHLGELHAILRRQEDFYPFLK
DNREKIEKILTFRIPYYVGPLARGNSRFAWMTRKSEETITPWNFEEVVDKGASAQSFIERMTNF
DKNLPNEKVLPKHSLLYEYFTVYNELTKVKYVTEGMRKPAFLSGEQKKAIVDLLFKTNRKVTVK
QLKEDYFKKIECFDSVEISGVEDRFNASLGTYHDLLKIIKDKDFLDNEENEDILEDIVLTLTLFEDR
EMIEERLKTYAHLFDDKVMKQLKRRRYTGWGRLSRKLINGIRDKQSGKTILDFLKSDGFANRNF
MQLIHDDSLTFKEDIQKAQVSGQGDSLHEHIANLAGSPAIKKGILQTVKVVDELVKVMGRHKPE
NIVIEMARENQTTQKGQKNSRERMKRIEEGIKELGSQILKEHPVENTQLQNEKLYLYYLQNGRD
MYVDQELDINRLSDYDVDHIVPQSFLKDDSIDNKVLTRSDKNRGKSDNVPSEEVVKKMKNYWR
QLLNAKLITQRKFDNLTKAERGGLSELDKAGFIKRQLVETRQITKHVAQILDSRMNTKYDENDKL

IREVKVITLKSKLVSDFRKDFQFYKVREINNYHHAHDAYLNAVVGTALIKKYPKLESEFVYGDYK
VYDVRKMIAKSEQEIGKATAKYFFYSNIMNFFKTEITLANGEIRKRPLIETNGETGEIVWDKGRDF
ATVRKVLSMPQVNIVKKTEVQTGGFSKESIRPKRNSDKLIARKKDWDPKKYGGFLWPTVAYSV
LVVAKVEKGKSKKLKSVKELLGITIMERSSFEKNPIDFLEAKGYKEVKKDLIIKLPKYSLFELENGR
KRMLASAKQLQKGNELALPSKYVNFLYLASHYEKLKGSPEDNEQKQLFVEQHKHYLDEIIEQIS
EFSKRVILADANLDKVLSAYNKHRDKPIREQAENIIHLFTLTRLGAPRAFKYFDTTIDPKQYRSTK
EVLDATLIHQSITGLYETRIDLSQLGGDSGGSGGSGGSTNLSDIIEKETGKQLVIQESILMLPEEV
EEVIGNKPESDILVHTAYDESTDENVMLLTSDAPEYKPWALVIQDSNGENKIKMLSGGSGGSG
GSTNLSDIIEKETGKQLVIQESILMLPEEVEEVIGNKPESDILVHTAYDESTDENVMLLTSDAPEY
KPWALVIQDSNGENKIKMLSGGSKRTADGSEFEPKKKRKVGGGGSGATNFSLLKQAGDVEEN
PGPMVSKGEELFTGVVPILVELDGDVNGHKFSVSGEGEGDATYGKLTLKFICTTGKLPVPWPTL
VTTLTYGVQCFSRYPDHMKQHDFFKSAMPEGYVQERTIFFKDDGNYKTRAEVKFEGDTLVNRI
ELKGIDFKEDGNILGHKLEYNYNSHNVYIMADKQKNGIKVNFKIRHNIEDGSVQLADHYQQNTPI
GDGPVLLPDNHYLSTQSALSKDPNEKRDHMVLLEFVTAAGITLGMDELYK*

N-SpRY ABE8e

MKRTADGSEFESPKKKRKVSEVEFSHEYWMRHALTLAKRARDEREVPVGAVLVLNNRVIGEG
WNRAIGLHDPTAHAEIMALRQGGLVMQNYRLIDATLYVTFEPCVMCAGAMIHSRIGRVVFGVR
NSKRGAAGSLMNVLNYPGMNHRVEITEGILADECAALLCDFYRMPRQVFNAQKKAQSSINSGG
SSGGSSGSETPGTSESATPESSGGSSGGSDKKYSIGLAIGTNSVGWAVITDEYKVPSKKFKVL
GNTDRHSIKKNLIGALLFDSGETAEATRLKRTARRRYTRRKNRICYLQEIFSNEMAKVDDSFFHR
LEESFLVEEDKKHERHPIFGNIVDEVAYHEKYPTIYHLRKKLVDSTDKADLRLIYLALAHMIKFRG
HFLIEGDLNPDNSDVDKLFIQLVQTYNQLFEENPINASGVDAKAILSARLSKSRRLENLIAQLPGE
KKNGLFGNLIALSLGLTPNFKSNFDLAEDAKLQLSKDTYDDDLDNLLAQIGDQYADLFLAAKNLS
DAILLSDILRVNTEITKAPLSASMIKRYDEHHQDLTLLKALVRQQLPEKYKEIFFDQSKNGYAGYI
DGGASQEEFYKFIKPILEKMDGTEELLVKLNREDLLRKQRTFDNGSIPHQIHLGELHAILRRQED
FYPFLKDNREKIEKILTFRIPYYVGPLARGNSRFAWMTRKSEETITPWNFEEVVDKGASAQSFIE
RMTNFDKNLPNEKVLPKHSLLYEYFTVYNELTKVKYVTEGMRKPAFLSGEQKKAIVDLLFKTNR
KVTVKQLKEDYFKKIECFDSVEISGVEDRFNASLGTYHDLLKIIKDKDFLDNEENEDILEDIVLTLT
LFEDREMIEERLKTYAHLFDDKVMKQLKRRRYTGWGRLSRKLINGIRDKQSGKTILDFLKSDGF
ANRNFMQLIHDDSLTFKEDIQKAQVCLAGDTLITLADGRRVPIRELVSQQNFSVWALNPQTYRL
ERARVSRAFCTGIKPVYRLTTRLGRSIRATANHRFLTPQGWKRVDELQPGDYLALPRRIPTAS*

C-SpRY ABE

MAAACPELRQLAQSDVYWDPIVSIEPDGVEEVFDLTVPGPHNFVANDIIAHNSGQGDSLHEHIA
NLAGSPAIKKGILQTVKVVDELVKVMGRHKPENIVIEMARENQTTQKGQKNSRERMKRIEEGIK
ELGSQILKEHPVENTQLQNEKLYLYYLQNGRDMYVDQELDINRLSDYDVDHIVPQSFLKDDSID
NKVLTRSDKNRGKSDNVPSEEVVKKMKNYWRQLLNAKLITQRKFDNLTKAERGGLSELDKAGF
IKRQLVETRQITKHVAQILDSRMNTKYDENDKLIREVKVITLKSKLVSDFRKDFQFYKVREINNYH
HAHDAYLNAVVGTALIKKYPKLESEFVYGDYKVYDVRKMIAKSEQEIGKATAKYFFYSNIMNFFK
TEITLANGEIRKRPLIETNGETGEIVWDKGRDFATVRKVLSMPQVNIVKKTEVQTGGFSKESILP
KRNSDKLIARKKDWDPKKYGGFLWPTVAYSVLVVAKVEKGKSKKLKSVKELLGITIMERSSFEK
NPIDFLEAKGYKEVKKDLIIKLPKYSLFELENGRKRMLASAKQLQKGNELALPSKYVNFLYLASH
YEKLKGSPEDNEQKQLFVEQHKHYLDEIIEQISEFSKRVILADANLDKVLSAYNKHRDKPIREQA
ENIIHLFTLTRLGAPRAFKYFDTTIDPKQYRSTKEVLDATLIHQSITGLYETRIDLSQLGGDSGGS
GGSGGSTNLSDIIEKETGKQLVIQESILMLPEEVEEVIGNKPESDILVHTAYDESTDENVMLLTSD
APEYKPWALVIQDSNGENKIKMLSGGSKRTADGSEFEPKKKRKV*

N-WT ABE8e

MKRTADGSEFESPKKKRKVSEVEFSHEYWMRHALTLAKRARDEREVPVGAVLVLNNRVIGEG
WNRAIGLHDPTAHAEIMALRQGGLVMQNYRLIDATLYVTFEPCVMCAGAMIHSRIGRVVFGVR
NSKRGAAGSLMNVLNYPGMNHRVEITEGILADECAALLCDFYRMPRQVFNAQKKAQSSINSGG
SSGGSSGSETPGTSESATPESSGGSSGGSDKKYSIGLAIGTNSVGWAVITDEYKVPSKKFKVL
GNTDRHSIKKNLIGALLFDSGETAEATRLKRTARRRYTRRKNRICYLQEIFSNEMAKVDDSFFHR
LEESFLVEEDKKHERHPIFGNIVDEVAYHEKYPTIYHLRKKLVDSTDKADLRLIYLALAHMIKFRG
HFLIEGDLNPDNSDVDKLFIQLVQTYNQLFEENPINASGVDAKAILSARLSKSRRLENLIAQLPGE
KKNGLFGNLIALSLGLTPNFKSNFDLAEDAKLQLSKDTYDDDLDNLLAQIGDQYADLFLAAKNLS
DAILLSDILRVNTEITKAPLSASMIKRYDEHHQDLTLLKALVRQQLPEKYKEIFFDQSKNGYAGYI
DGGASQEEFYKFIKPILEKMDGTEELLVKLNREDLLRKQRTFDNGSIPHQIHLGELHAILRRQED
FYPFLKDNREKIEKILTFRIPYYVGPLARGNSRFAWMTRKSEETITPWNFEEVVDKGASAQSFIE
RMTNFDKNLPNEKVLPKHSLLYEYFTVYNELTKVKYVTEGMRKPAFLSGEQKKAIVDLLFKTNR
KVTVKQLKEDYFKKIECFDSVEISGVEDRFNASLGTYHDLLKIIKDKDFLDNEENEDILEDIVLTLT
LFEDREMIEERLKTYAHLFDDKVMKQLKRRRYTGWGRLSRKLINGIRDKQSGKTILDFLKSDGF
ANRNFMQLIHDDSLTFKEDIQKAQVCLAGDTLITLADGRRVPIRELVSQQNFSVWALNPQTYRL
ERARVSRAFCTGIKPVYRLTTRLGRSIRATANHRFLTPQGWKRVDELQPGDYLALPRRIPTAS*

C-WT ABE

MAAACPELRQLAQSDVYWDPIVSIEPDGVEEVFDLTVPGPHNFVANDIIAHNSGQGDSLHEHIA
NLAGSPAIKKGILQTVKVVDELVKVMGRHKPENIVIEMARENQTTQKGQKNSRERMKRIEEGIK
ELGSQILKEHPVENTQLQNEKLYLYYLQNGRDMYVDQELDINRLSDYDVDHIVPQSFLKDDSID
NKVLTRSDKNRGKSDNVPSEEVVKKMKNYWRQLLNAKLITQRKFDNLTKAERGGLSELDKAGF
IKRQLVETRQITKHVAQILDSRMNTKYDENDKLIREVKVITLKSKLVSDFRKDFQFYKVREINNYH
HAHDAYLNAVVGTALIKKYPKLESEFVYGDYKVYDVRKMIAKSEQEIGKATAKYFFYSNIMNFFK
TEITLANGEIRKRPLIETNGETGEIVWDKGRDFATVRKVLSMPQVNIVKKTEVQTGGFSKESILP
KRNSDKLIARKKDWDPKKYGGFDSPTVAYSVLVVAKVEKGKSKKLKSVKELLGITIMERSSFEK
NPIDFLEAKGYKEVKKDLIIKLPKYSLFELENGRKRMLASAGELQKGNELALPSKYVNFLYLASH
YEKLKGSPEDNEQKQLFVEQHKHYLDEIIEQISEFSKRVILADANLDKVLSAYNKHRDKPIREQA
ENIIHLFTLTNLGAPAAFKYFDTTIDRKRYTSTKEVLDATLIHQSITGLYETRIDLSQLGGDSGGSG
GSGGSTNLSDIIEKETGKQLVIQESILMLPEEVEEVIGNKPESDILVHTAYDESTDENVMLLTSDA
PEYKPWALVIQDSNGENKIKMLSGGSKRTADGSEFEPKKKRKV*

N-SpRY ABE7.10

MKRTADGSEFESPKKKRKVSEVEFSHEYWMRHALTLAKRAWDEREVPVGAVLVHNNRVIGEG
WNRPIGRHDPTAHAEIMALRQGGLVMQNYRLIDATLYVTLEPCVMCAGAMIHSRIGRVVFGAR
DAKTGAAGSLMDVLHHPGMNHRVEITEGILADECAALLSDFFRMRRQEIKAQKKAQSSTDSGG
SSGGSSGSETPGTSESATPESSGGSSGGSSEVEFSHEYWMRHALTLAKRARDEREVPVGAV
LVLNNRVIGEGWNRAIGLHDPTAHAEIMALRQGGLVMQNYRLIDATLYVTFEPCVMCAGAMIHS
RIGRVVFGVRNAKTGAAGSLMDVLHYPGMNHRVEITEGILADECAALLCYFFRMPRQVFNAQK
KAQSSTDASGGSSGGSSGSETPGTSESATPESSGGSSGGSDKKYSIGLAIGTNSVGWAVITDE
YKVPSKKFKVLGNTDRHSIKKNLIGALLFDSGETAEATRLKRTARRRYTRRKNRICYLQEIFSNE
MAKVDDSFFHRLEESFLVEEDKKHERHPIFGNIVDEVAYHEKYPTIYHLRKKLVDSTDKADLRLI
YLALAHMIKFRGHFLIEGDLNPDNSDVDKLFIQLVQTYNQLFEENPINASGVDAKAILSARLSKSR
RLENLIAQLPGEKKNGLFGNLIALSLGLTPNFKSNFDLAEDAKLQLSKDTYDDDLDNLLAQIGDQ
YADLFLAAKNLSDAILLSDILRVNTEITKAPLSASMIKRYDEHHQDLTLLKALVRQQLPEKYKEIFF
DQSKNGYAGYIDGGASQEEFYKFIKPILEKMDGTEELLVKLNREDLLRKQRTFDNGSIPHQIHLG
ELHAILRRQEDFYPFLKDNREKIEKILTFRIPYYVGPLARGNSRFAWMTRKSEETITPWNFEEVV
DKGASAQSFIERMTNFDKNLPNEKVLPKHSLLYEYFTVYNELTKVKYVTEGMRKPAFLSGEQK
KAIVDLLFKTNRKVTVKQLKEDYFKKIECFDSVEISGVEDRFNASLGTYHDLLKIIKDKDFLDNEE

NEDILEDIVLTLTLFEDREMIEERLKTYAHLFDDKVMKQLKRRRYTGWGRLSRKLINGIRDKQSG
KTILDFLKSDGFANRNFMQLIHDDSLTFKEDIQKAQVCLAGDTLITLADGRRVPIRELVSQQNFS
VWALNPQTYRLERARVSRAFCTGIKPVYRLTTRLGRSIRATANHRFLTPQGWKRVDELQPGDY
LALPRRIPTAS*

N-SpRY ABE8.20m

MKRTADGSEFESPKKKRKVSEVEFSHEYWMRHALTLAKRARDEREVPVGAVLVLNNRVIGEG
WNRAIGLHDPTAHAEIMALRQGGLVMQNYRLYDATLYSTFEPCVMCAGAMIHSRIGRVVFGVR
NAKTGAAGSLMDVLHHPGMNHRVEITEGILADECAALLCRFFRMPRRVFNAQKKAQSSTDSG
GSSGGSSGSETPGTSESATPESSGGSSGGSDKKYSIGLAIGTNSVGWAVITDEYKVPSKKFKV
LGNTDRHSIKKNLIGALLFDSGETAEATRLKRTARRRYTRRKNRICYLQEIFSNEMAKVDDSFFH
RLEESFLVEEDKKHERHPIFGNIVDEVAYHEKYPTIYHLRKKLVDSTDKADLRLIYLALAHMIKFR
GHFLIEGDLNPDNSDVDKLFIQLVQTYNQLFEENPINASGVDAKAILSARLSKSRRLENLIAQLPG
EKKNGLFGNLIALSLGLTPNFKSNFDLAEDAKLQLSKDTYDDDLDNLLAQIGDQYADLFLAAKNL
SDAILLSDILRVNTEITKAPLSASMIKRYDEHHQDLTLLKALVRQQLPEKYKEIFFDQSKNGYAGY
IDGGASQEEFYKFIKPILEKMDGTEELLVKLNREDLLRKQRTFDNGSIPHQIHLGELHAILRRQED
FYPFLKDNREKIEKILTFRIPYYVGPLARGNSRFAWMTRKSEETITPWNFEEVVDKGASAQSFIE
RMTNFDKNLPNEKVLPKHSLLYEYFTVYNELTKVKYVTEGMRKPAFLSGEQKKAIVDLLFKTNR
KVTVKQLKEDYFKKIECFDSVEISGVEDRFNASLGTYHDLLKIIKDKDFLDNEENEDILEDIVLTLT
LFEDREMIEERLKTYAHLFDDKVMKQLKRRRYTGWGRLSRKLINGIRDKQSGKTILDFLKSDGF
ANRNFMQLIHDDSLTFKEDIQKAQVSGQGDSLHEHIANLAGSPAIKKGILQTVKVVDELVKVMG
RHKPENIVIEMARENQTTQKGQKNSRERMKRIEEGIKELGSQILKEHPVENTQLQNEKLYLYYL
QNGRDMYVDQELDINRLSDYDVDHIVPQSFLKDDSIDNKVLTRSDKNRGKSDNVPSEEVVKKM
KNYWRQLLNAKLITQRKFDNLTKAERGGLSELDKAGFIKRQLVETRQITKHVAQILDSRMNTKY
DENDKLIREVKVITLKSKLVSDFRKDFQFYKVREINNYHHAHDAYLNAVVGTALIKKYPKLESEFV
YGDYKVYDVRKMIAKSEQEIGKATAKYFFYSNIMNFFKTEITLANGEIRKRPLIETNGETGEIVWD
KGRDFATVRKVLSMPQVNIVKKTEVQTGGFSKESILPKRNSDKLIARKKDWDPKKYGGFLWPT
VAYSVLVVAKVEKGKSKKLKSVKELLGITIMERSSFEKNPIDFLEAKGYKEVKKDLIIKLPKYSLFE
LENGRKRMLASAKQLQKGNELALPSKYVNFLYLASHYEKLKGSPEDNEQKQLFVEQHKHYLDE
IIEQISEFSKRVILADANLDKVLSAYNKHRDKPIREQAENIIHLFTLTRLGAPRAFKYFDTTIDPKQY
RSTKEVLDATLIHQSITGLYETRIDLSQLGGDSGGSKRTADGSEFEPKKKRKV*

N-SpRY ABE9

MKRTADGSEFESPKKKRKVSEVEFSHEYWMRHALTLAKRARDEREVPVGAVLVLNNRVIGEG
WNRAIGLHDPTAHAEIMALRQGGLVMQNYRLIDATLYVTFEPCVMCAGAMIHSRIGRVVFGVR
QSKRGAAGSLMNVLNYPGMNHRVEITEGILADECAALTCDFYRMPRQVFNAQKKAQSSINSG
GSSGGSSGSETPGTSESATPESSGGSSGGSDKKYSIGLAIGTNSVGWAVITDEYKVPSKKFKV
LGNTDRHSIKKNLIGALLFDSGETAEATRLKRTARRRYTRRKNRICYLQEIFSNEMAKVDDSFFH
RLEESFLVEEDKKHERHPIFGNIVDEVAYHEKYPTIYHLRKKLVDSTDKADLRLIYLALAHMIKFR
GHFLIEGDLNPDNSDVDKLFIQLVQTYNQLFEENPINASGVDAKAILSARLSKSRRLENLIAQLPG
EKKNGLFGNLIALSLGLTPNFKSNFDLAEDAKLQLSKDTYDDDLDNLLAQIGDQYADLFLAAKNL
SDAILLSDILRVNTEITKAPLSASMIKRYDEHHQDLTLLKALVRQQLPEKYKEIFFDQSKNGYAGY
IDGGASQEEFYKFIKPILEKMDGTEELLVKLNREDLLRKQRTFDNGSIPHQIHLGELHAILRRQED
FYPFLKDNREKIEKILTFRIPYYVGPLARGNSRFAWMTRKSEETITPWNFEEVVDKGASAQSFIE
RMTNFDKNLPNEKVLPKHSLLYEYFTVYNELTKVKYVTEGMRKPAFLSGEQKKAIVDLLFKTNR
KVTVKQLKEDYFKKIECFDSVEISGVEDRFNASLGTYHDLLKIIKDKDFLDNEENEDILEDIVLTLT
LFEDREMIEERLKTYAHLFDDKVMKQLKRRRYTGWGRLSRKLINGIRDKQSGKTILDFLKSDGF
ANRNFMQLIHDDSLTFKEDIQKAQVCLAGDTLITLADGRRVPIRELVSQQNFSVWALNPQTYRL
ERARVSRAFCTGIKPVYRLTTRLGRSIRATANHRFLTPQGWKRVDELQPGDYLALPRRIPTAS*

N-SpRY CBE4max

MGKPIPNPLLGLDSTKRTADGSEFESPKKKRKVSSETGPVAVDPTLRRRIEPHEFEVFFDPREL
RKETCLLYEINWGGRHSIWRHTSQNTNKHVEVNFIEKFTTERYFCPNTRCSITWFLSWSPCGE
CSRAITEFLSRYPHVTLFIYIARLYHHADPRNRQGLRDLISSGVTIQIMTEQESGYCWRNFVNYS
PSNEAHWPRYPHLWVRLYVLELYCIILGLPPCLNILRRKQPQLTFFTIALQSCHYQRLPPHILWA
TGLKSGGSSGGSSGSETPGTSESATPESSGGSSGGSDKKYSIGLAIGTNSVGWAVITDEYKVP
SKKFKVLGNTDRHSIKKNLIGALLFDSGETAEATRLKRTARRRYTRRKNRICYLQEIFSNEMAKV
DDSFFHRLEESFLVEEDKKHERHPIFGNIVDEVAYHEKYPTIYHLRKKLVDSTDKADLRLIYLALA
HMIKFRGHFLIEGDLNPDNSDVDKLFIQLVQTYNQLFEENPINASGVDAKAILSARLSKSRRLEN
LIAQLPGEKKNGLFGNLIALSLGLTPNFKSNFDLAEDAKLQLSKDTYDDDLDNLLAQIGDQYADL
FLAAKNLSDAILLSDILRVNTEITKAPLSASMIKRYDEHHQDLTLLKALVRQQLPEKYKEIFFDQSK
NGYAGYIDGGASQEEFYKFIKPILEKMDGTEELLVKLNREDLLRKQRTFDNGSIPHQIHLGELHAI
LRRQEDFYPFLKDNREKIEKILTFRIPYYVGPLARGNSRFAWMTRKSEETITPWNFEEVVDKGA
SAQSFIERMTNFDKNLPNEKVLPKHSLLYEYFTVYNELTKVKYVTEGMRKPAFLSGEQKKAIVD
LLFKTNRKVTVKQLKEDYFKKIECFDSVEISGVEDRFNASLGTYHDLLKIIKDKDFLDNEENEDIL
EDIVLTLTLFEDREMIEERLKTYAHLFDDKVMKQLKRRRYTGWGRLSRKLINGIRDKQSGKTILD
FLKSDGFANRNFMQLIHDDSLTFKEDIQKAQVCLAGDTLITLADGRRVPIRELVSQQNFSVWAL
NPQTYRLERARVSRAFCTGIKPVYRLTTRLGRSIRATANHRFLTPQGWKRVDELQPGDYLALP
RRIPTAS*

C-SpRY CBE

MAAACPELRQLAQSDVYWDPIVSIEPDGVEEVFDLTVPGPHNFVANDIIAHNSGQGDSLHEHIA
NLAGSPAIKKGILQTVKVVDELVKVMGRHKPENIVIEMARENQTTQKGQKNSRERMKRIEEGIK
ELGSQILKEHPVENTQLQNEKLYLYYLQNGRDMYVDQELDINRLSDYDVDHIVPQSFLKDDSID
NKVLTRSDKNRGKSDNVPSEEVVKKMKNYWRQLLNAKLITQRKFDNLTKAERGGLSELDKAGF
IKRQLVETRQITKHVAQILDSRMNTKYDENDKLIREVKVITLKSKLVSDFRKDFQFYKVREINNYH
HAHDAYLNAVVGTALIKKYPKLESEFVYGDYKVYDVRKMIAKSEQEIGKATAKYFFYSNIMNFFK
TEITLANGEIRKRPLIETNGETGEIVWDKGRDFATVRKVLSMPQVNIVKKTEVQTGGFSKESIRP
KRNSDKLIARKKDWDPKKYGGFLWPTVAYSVLVVAKVEKGKSKKLKSVKELLGITIMERSSFEK
NPIDFLEAKGYKEVKKDLIIKLPKYSLFELENGRKRMLASAKQLQKGNELALPSKYVNFLYLASH
YEKLKGSPEDNEQKQLFVEQHKHYLDEIIEQISEFSKRVILADANLDKVLSAYNKHRDKPIREQA
ENIIHLFTLTRLGAPRAFKYFDTTIDPKQYRSTKEVLDATLIHQSITGLYETRIDLSQLGGDSGGS
GGSGGSTNLSDIIEKETGKQLVIQESILMLPEEVEEVIGNKPESDILVHTAYDESTDENVMLLTSD
APEYKPWALVIQDSNGENKIKMLSGGSGGSGGSTNLSDIIEKETGKQLVIQESILMLPEEVEEVI
GNKPESDILVHTAYDESTDENVMLLTSDAPEYKPWALVIQDSNGENKIKMLYPYDVPDYAYPYD
VPDYAYPYDVPDYASGGSPKKKRKV*

N-WT CBE4max

MGKPIPNPLLGLDSTKRTADGSEFESPKKKRKVSSETGPVAVDPTLRRRIEPHEFEVFFDPREL
RKETCLLYEINWGGRHSIWRHTSQNTNKHVEVNFIEKFTTERYFCPNTRCSITWFLSWSPCGE
CSRAITEFLSRYPHVTLFIYIARLYHHADPRNRQGLRDLISSGVTIQIMTEQESGYCWRNFVNYS
PSNEAHWPRYPHLWVRLYVLELYCIILGLPPCLNILRRKQPQLTFFTIALQSCHYQRLPPHILWA
TGLKSGGSSGGSSGSETPGTSESATPESSGGSSGGSDKKYSIGLAIGTNSVGWAVITDEYKVP
SKKFKVLGNTDRHSIKKNLIGALLFDSGETAEATRLKRTARRRYTRRKNRICYLQEIFSNEMAKV
DDSFFHRLEESFLVEEDKKHERHPIFGNIVDEVAYHEKYPTIYHLRKKLVDSTDKADLRLIYLALA
HMIKFRGHFLIEGDLNPDNSDVDKLFIQLVQTYNQLFEENPINASGVDAKAILSARLSKSRRLEN
LIAQLPGEKKNGLFGNLIALSLGLTPNFKSNFDLAEDAKLQLSKDTYDDDLDNLLAQIGDQYADL
FLAAKNLSDAILLSDILRVNTEITKAPLSASMIKRYDEHHQDLTLLKALVRQQLPEKYKEIFFDQSK

NGYAGYIDGGASQEEFYKFIKPILEKMDGTEELLVKLNREDLLRKQRTFDNGSIPHQIHLGELHAI
LRRQEDFYPFLKDNREKIEKILTFRIPYYVGPLARGNSRFAWMTRKSEETITPWNFEEVVDKGA
SAQSFIERMTNFDKNLPNEKVLPKHSLLYEYFTVYNELTKVKYVTEGMRKPAFLSGEQKKAIVD
LLFKTNRKVTVKQLKEDYFKKIECFDSVEISGVEDRFNASLGTYHDLLKIIKDKDFLDNEENEDIL
EDIVLTLTLFEDREMIEERLKTYAHLFDDKVMKQLKRRRYTGWGRLSRKLINGIRDKQSGKTILD
FLKSDGFANRNFMQLIHDDSLTFKEDIQKAQVCLAGDTLITLADGRRVPIRELVSQQNFSVWAL
NPQTYRLERARVSRAFCTGIKPVYRLTTRLGRSIRATANHRFLTPQGWKRVDELQPGDYLALP
RRIPTAS*

C-WT CBE4max

MAAACPELRQLAQSDVYWDPIVSIEPDGVEEVFDLTVPGPHNFVANDIIAHNSGQGDSLHEHIA
NLAGSPAIKKGILQTVKVVDELVKVMGRHKPENIVIEMARENQTTQKGQKNSRERMKRIEEGIK
ELGSQILKEHPVENTQLQNEKLYLYYLQNGRDMYVDQELDINRLSDYDVDHIVPQSFLKDDSID
NKVLTRSDKNRGKSDNVPSEEVVKKMKNYWRQLLNAKLITQRKFDNLTKAERGGLSELDKAGF
IKRQLVETRQITKHVAQILDSRMNTKYDENDKLIREVKVITLKSKLVSDFRKDFQFYKVREINNYH
HAHDAYLNAVVGTALIKKYPKLESEFVYGDYKVYDVRKMIAKSEQEIGKATAKYFFYSNIMNFFK
TEITLANGEIRKRPLIETNGETGEIVWDKGRDFATVRKVLSMPQVNIVKKTEVQTGGFSKESILP
KRNSDKLIARKKDWDPKKYGGFDSPTVAYSVLVVAKVEKGKSKKLKSVKELLGITIMERSSFEK
NPIDFLEAKGYKEVKKDLIIKLPKYSLFELENGRKRMLASAGELQKGNELALPSKYVNFLYLASH
YEKLKGSPEDNEQKQLFVEQHKHYLDEIIEQISEFSKRVILADANLDKVLSAYNKHRDKPIREQA
ENIIHLFTLTNLGAPAAFKYFDTTIDRKRYTSTKEVLDATLIHQSITGLYETRIDLSQLGGDSGGST
NLSDIIEKETGKQLVIQESILMLPEEVEEVIGNKPESDILVHTAYDESTDENVMLLTSDAPEYKPW
ALVIQDSNGENKIKMLSYPYDVPDYAYPYDVPDYAYPYDVPDYASGGSPKKKRKV*

N-SpRY PpAPOBEC1(H122A)

MGKPIPNPLLGLDSTKRTADGSEFESPKKKRKVTSEKGPSTGDPTLRRRIESWEFDVFYDPRE
LRKETCLLYEIKWGMSRKIWRSSGKNTTNHVEVNFIKKFTSERRFHSSISCSITWFLSWSPCWE
CSQAIREFLSQHPGVTLVIYVARLFWAMDQRNRQGLRDLVNSGVTIQIMRASEYYHCWRNFVN
YPPGDEAHWPQYPPLWMMLYALELHCIILSLPPCLKISRRWQNHLAFFRLHLQNCHYQTIPPHI
LLATGLIHPSVTWRLKSGGSSGGSSGSETPGTSESATPESSGGSSGGSDKKYSIGLAIGTNSV
GWAVITDEYKVPSKKFKVLGNTDRHSIKKNLIGALLFDSGETAEATRLKRTARRRYTRRKNRICY
LQEIFSNEMAKVDDSFFHRLEESFLVEEDKKHERHPIFGNIVDEVAYHEKYPTIYHLRKKLVDST
DKADLRLIYLALAHMIKFRGHFLIEGDLNPDNSDVDKLFIQLVQTYNQLFEENPINASGVDAKAIL
SARLSKSRRLENLIAQLPGEKKNGLFGNLIALSLGLTPNFKSNFDLAEDAKLQLSKDTYDDDLDN
LLAQIGDQYADLFLAAKNLSDAILLSDILRVNTEITKAPLSASMIKRYDEHHQDLTLLKALVRQQLP
EKYKEIFFDQSKNGYAGYIDGGASQEEFYKFIKPILEKMDGTEELLVKLNREDLLRKQRTFDNGS
IPHQIHLGELHAILRRQEDFYPFLKDNREKIEKILTFRIPYYVGPLARGNSRFAWMTRKSEETITP
WNFEEVVDKGASAQSFIERMTNFDKNLPNEKVLPKHSLLYEYFTVYNELTKVKYVTEGMRKPA
FLSGEQKKAIVDLLFKTNRKVTVKQLKEDYFKKIECFDSVEISGVEDRFNASLGTYHDLLKIIKDK
DFLDNEENEDILEDIVLTLTLFEDREMIEERLKTYAHLFDDKVMKQLKRRRYTGWGRLSRKLING
IRDKQSGKTILDFLKSDGFANRNFMQLIHDDSLTFKEDIQKAQVCLAGDTLITLADGRRVPIRELV
SQQNFSVWALNPQTYRLERARVSRAFCTGIKPVYRLTTRLGRSIRATANHRFLTPQGWKRVDE
LQPGDYLALPRRIPTAS*

N-SpRY evoFERNY

MGKPIPNPLLGLDSTKRTADGSEFESPKKKRKVSFERNYDPRELRKETYLLYEIKWGKSGKLW
RHWCQNNRTQHAEVYFLENIFNARRFNPSTHCSITWYLSWSPCAECSQKIVDFLKEHPNVNLE
IYVARLYYPENERNRQGLRDLVNSGVTIRIMDLPDYNYCWKTFVSDQGGDEDYWPGHFAPWIK

QYSLKLSGGSSGGSSGSETPGTSESATPESSGGSSGGSDKKYSIGLAIGTNSVGWAVITDEYK
VPSKKFKVLGNTDRHSIKKNLIGALLFDSGETAEATRLKRTARRRYTRRKNRICYLQEIFSNEMA
KVDDSFFHRLEESFLVEEDKKHERHPIFGNIVDEVAYHEKYPTIYHLRKKLVDSTDKADLRLIYLA
LAHMIKFRGHFLIEGDLNPDNSDVDKLFIQLVQTYNQLFEENPINASGVDAKAILSARLSKSRRLE
NLIAQLPGEKKNGLFGNLIALSLGLTPNFKSNFDLAEDAKLQLSKDTYDDDLDNLLAQIGDQYAD
LFLAAKNLSDAILLSDILRVNTEITKAPLSASMIKRYDEHHQDLTLLKALVRQQLPEKYKEIFFDQS
KNGYAGYIDGGASQEEFYKFIKPILEKMDGTEELLVKLNREDLLRKQRTFDNGSIPHQIHLGELH
AILRRQEDFYPFLKDNREKIEKILTFRIPYYVGPLARGNSRFAWMTRKSEETITPWNFEEVVDKG
ASAQSFIERMTNFDKNLPNEKVLPKHSLLYEYFTVYNELTKVKYVTEGMRKPAFLSGEQKKAIV
DLLFKTNRKVTVKQLKEDYFKKIECFDSVEISGVEDRFNASLGTYHDLLKIIKDKDFLDNEENEDI
LEDIVLTLTLFEDREMIEERLKTYAHLFDDKVMKQLKRRRYTGWGRLSRKLINGIRDKQSGKTIL
DFLKSDGFANRNFMQLIHDDSLTFKEDIQKAQVCLAGDTLITLADGRRVPIRELVSQQNFSVWA
LNPQTYRLERARVSRAFCTGIKPVYRLTTRLGRSIRATANHRFLTPQGWKRVDELQPGDYLALP
RRIPTAS*

N-SpRY SsAPOBEC3B(R54Q)

MGKPIPNPLLGLDSTKRTADGSEFESPKKKRKVDPQRLRQWPGPGPASRGGYGQRPRIRNPE
EWFHELSPRTFSFHFRNLRFASGQNRSYICCQVEGKNCFFQGIFQNQVPPDPPCHAELCFLS
WFQSWGLSPDEHYYVTWFISWSPCCECAAKVAQFLEENRNVSLSLSAARLYYFWKSESREGL
RRLSDLGAQVGIMSFQDFQHCWNNFVHNLGMPFQPWKKLHKNYQRLVTELKQILREEPATYG
SPQAQGKVRIGSTAAGLRHSHSHTRSEAHLRPNHSSRQHRILNPPREARARTCVLVDASWICY
RSGGSSGGSSGSETPGTSESATPESSGGSSGGSDKKYSIGLAIGTNSVGWAVITDEYKVPSKK
FKVLGNTDRHSIKKNLIGALLFDSGETAEATRLKRTARRRYTRRKNRICYLQEIFSNEMAKVDDS
FFHRLEESFLVEEDKKHERHPIFGNIVDEVAYHEKYPTIYHLRKKLVDSTDKADLRLIYLALAHMI
KFRGHFLIEGDLNPDNSDVDKLFIQLVQTYNQLFEENPINASGVDAKAILSARLSKSRRLENLIAQ
LPGEKKNGLFGNLIALSLGLTPNFKSNFDLAEDAKLQLSKDTYDDDLDNLLAQIGDQYADLFLAA
KNLSDAILLSDILRVNTEITKAPLSASMIKRYDEHHQDLTLLKALVRQQLPEKYKEIFFDQSKNGY
AGYIDGGASQEEFYKFIKPILEKMDGTEELLVKLNREDLLRKQRTFDNGSIPHQIHLGELHAILRR
QEDFYPFLKDNREKIEKILTFRIPYYVGPLARGNSRFAWMTRKSEETITPWNFEEVVDKGASAQ
SFIERMTNFDKNLPNEKVLPKHSLLYEYFTVYNELTKVKYVTEGMRKPAFLSGEQKKAIVDLLFK
TNRKVTVKQLKEDYFKKIECFDSVEISGVEDRFNASLGTYHDLLKIIKDKDFLDNEENEDILEDIVL
TLTLFEDREMIEERLKTYAHLFDDKVMKQLKRRRYTGWGRLSRKLINGIRDKQSGKTILDFLKSD
GFANRNFMQLIHDDSLTFKEDIQKAQVCLAGDTLITLADGRRVPIRELVSQQNFSVWALNPQTY
RLERARVSRAFCTGIKPVYRLTTRLGRSIRATANHRFLTPQGWKRVDELQPGDYLALPRRIPTA
S*

N-SpRY RrA3F(F130L)

MGKPIPNPLLGLDSTKRTADGSEFESPKKKRKVKPQIRDHRPNPMEAMYPHIFYFHFENLEKAY
GRNETWLCFTVEIIKQYLPVPWKKGVFRNQVDPETHCHAEKCFLSWFCNNTLSPKKNYQVTW
YTSWSPCPECAGEVAEFLAEHSNVKLTIYTARLYYLWDTDYQEGLRSLSEEGASVEIMDYEDF
QYCWENFVYDDGEPFKRWKGLKYNFQSLTRRLREILQSGGSSGGSSGSETPGTSESATPESS
GGSSGGSDKKYSIGLAIGTNSVGWAVITDEYKVPSKKFKVLGNTDRHSIKKNLIGALLFDSGETA
EATRLKRTARRRYTRRKNRICYLQEIFSNEMAKVDDSFFHRLEESFLVEEDKKHERHPIFGNIVD
EVAYHEKYPTIYHLRKKLVDSTDKADLRLIYLALAHMIKFRGHFLIEGDLNPDNSDVDKLFIQLVQ
TYNQLFEENPINASGVDAKAILSARLSKSRRLENLIAQLPGEKKNGLFGNLIALSLGLTPNFKSNF
DLAEDAKLQLSKDTYDDDLDNLLAQIGDQYADLFLAAKNLSDAILLSDILRVNTEITKAPLSASMIK
RYDEHHQDLTLLKALVRQQLPEKYKEIFFDQSKNGYAGYIDGGASQEEFYKFIKPILEKMDGTE
ELLVKLNREDLLRKQRTFDNGSIPHQIHLGELHAILRRQEDFYPFLKDNREKIEKILTFRIPYYVGP
LARGNSRFAWMTRKSEETITPWNFEEVVDKGASAQSFIERMTNFDKNLPNEKVLPKHSLLYEY

FTVYNELTKVKYVTEGMRKPAFLSGEQKKAIVDLLFKTNRKVTVKQLKEDYFKKIECFDSVEISG
VEDRFNASLGTYHDLLKIIKDKDFLDNEENEDILEDIVLTLTLFEDREMIEERLKTYAHLFDDKVM
KQLKRRRYTGWGRLSRKLINGIRDKQSGKTILDFLKSDGFANRNFMQLIHDDSLTFKEDIQKAQ
VCLAGDTLITLADGRRVPIRELVSQQNFSVWALNPQTYRLERARVSRAFCTGIKPVYRLTTRLG
RSIRATANHRFLTPQGWKRVDELQPGDYLALPRRIPTAS*

N-SpRY TadCBEd

MGKPIPNPLLGLDSTKRTADGSEFESPKKKRKVSEVEFSHEYWMRHALTLAKRARDERKAPVG
AVLVLNNRVIGEGWNRAIGLHDPTAHAEIIALRQGGLVMQNYRLIDATLYVTFEPCVMCAGAMIN
SRIGRVVFGVRNSKRGAAGSLMNVLNYPGMNHRVEITEGILADECAALLCDFYRMPRQVFNAQ
KKAQSSINSGGSSGGSSGSETPGTSESATPESSGGSSGGSDKKYSIGLAIGTNSVGWAVITDE
YKVPSKKFKVLGNTDRHSIKKNLIGALLFDSGETAEATRLKRTARRRYTRRKNRICYLQEIFSNE
MAKVDDSFFHRLEESFLVEEDKKHERHPIFGNIVDEVAYHEKYPTIYHLRKKLVDSTDKADLRLI
YLALAHMIKFRGHFLIEGDLNPDNSDVDKLFIQLVQTYNQLFEENPINASGVDAKAILSARLSKSR
RLENLIAQLPGEKKNGLFGNLIALSLGLTPNFKSNFDLAEDAKLQLSKDTYDDDLDNLLAQIGDQ
YADLFLAAKNLSDAILLSDILRVNTEITKAPLSASMIKRYDEHHQDLTLLKALVRQQLPEKYKEIFF
DQSKNGYAGYIDGGASQEEFYKFIKPILEKMDGTEELLVKLNREDLLRKQRTFDNGSIPHQIHLG
ELHAILRRQEDFYPFLKDNREKIEKILTFRIPYYVGPLARGNSRFAWMTRKSEETITPWNFEEVV
DKGASAQSFIERMTNFDKNLPNEKVLPKHSLLYEYFTVYNELTKVKYVTEGMRKPAFLSGEQK
KAIVDLLFKTNRKVTVKQLKEDYFKKIECFDSVEISGVEDRFNASLGTYHDLLKIIKDKDFLDNEE
NEDILEDIVLTLTLFEDREMIEERLKTYAHLFDDKVMKQLKRRRYTGWGRLSRKLINGIRDKQSG
KTILDFLKSDGFANRNFMQLIHDDSLTFKEDIQKAQVCLAGDTLITLADGRRVPIRELVSQQNFS
VWALNPQTYRLERARVSRAFCTGIKPVYRLTTRLGRSIRATANHRFLTPQGWKRVDELQPGDY
LALPRRIPTAS*

N-SpRY ABE8e (in vivo studies)

MGKPIPNPLLGLDSTKRTADGSEFESPKKKRKVSEVEFSHEYWMRHALTLAKRARDEREVPVG
AVLVLNNRVIGEGWNRAIGLHDPTAHAEIMALRQGGLVMQNYRLIDATLYVTFEPCVMCAGAMI
HSRIGRVVFGVRNSKRGAAGSLMNVLNYPGMNHRVEITEGILADECAALLCDFYRMPRQVFNA
QKKAQSSINSGGSSGGSSGSETPGTSESATPESSGGSSGGSDKKYSIGLAIGTNSVGWAVITD
EYKVPSKKFKVLGNTDRHSIKKNLIGALLFDSGETAEATRLKRTARRRYTRRKNRICYLQEIFSN
EMAKVDDSFFHRLEESFLVEEDKKHERHPIFGNIVDEVAYHEKYPTIYHLRKKLVDSTDKADLRL
IYLALAHMIKFRGHFLIEGDLNPDNSDVDKLFIQLVQTYNQLFEENPINASGVDAKAILSARLSKS
RRLENLIAQLPGEKKNGLFGNLIALSLGLTPNFKSNFDLAEDAKLQLSKDTYDDDLDNLLAQIGD
QYADLFLAAKNLSDAILLSDILRVNTEITKAPLSASMIKRYDEHHQDLTLLKALVRQQLPEKYKEIF
FDQSKNGYAGYIDGGASQEEFYKFIKPILEKMDGTEELLVKLNREDLLRKQRTFDNGSIPHQIHL
GELHAILRRQEDFYPFLKDNREKIEKILTFRIPYYVGPLARGNSRFAWMTRKSEETITPWNFEEV
VDKGASAQSFIERMTNFDKNLPNEKVLPKHSLLYEYFTVYNELTKVKYVTEGMRKPAFLSGEQ
KKAIVDLLFKTNRKVTVKQLKEDYFKKIECFDSVEISGVEDRFNASLGTYHDLLKIIKDKDFLDNE
ENEDILEDIVLTLTLFEDREMIEERLKTYAHLFDDKVMKQLKRRRYTGWGRLSRKLINGIRDKQS
GKTILDFLKSDGFANRNFMQLIHDDSLTFKEDIQKAQVCLAGDTLITLADGRRVPIRELVSQQNF
SVWALNPQTYRLERARVSRAFCTGIKPVYRLTTRLGRSIRATANHRFLTPQGWKRVDELQPGD
YLALPRRIPTAS*

C-SpRY ABE8e (in vivo studies)

MAAACPELRQLAQSDVYWDPIVSIEPDGVEEVFDLTVPGPHNFVANDIIAHNSGQGDSLHEHIA
NLAGSPAIKKGILQTVKVVDELVKVMGRHKPENIVIEMARENQTTQKGQKNSRERMKRIEEGIK
ELGSQILKEHPVENTQLQNEKLYLYYLQNGRDMYVDQELDINRLSDYDVDHIVPQSFLKDDSID

NKVLTRSDKNRGKSDNVPSEEVVKKMKNYWRQLLNAKLITQRKFDNLTKAERGGLSELDKAGF
IKRQLVETRQITKHVAQILDSRMNTKYDENDKLIREVKVITLKSKLVSDFRKDFQFYKVREINNYH
HAHDAYLNAVVGTALIKKYPKLESEFVYGDYKVYDVRKMIAKSEQEIGKATAKYFFYSNIMNFFK
TEITLANGEIRKRPLIETNGETGEIVWDKGRDFATVRKVLSMPQVNIVKKTEVQTGGFSKESILP
KRNSDKLIARKKDWDPKKYGGFLWPTVAYSVLVVAKVEKGKSKKLKSVKELLGITIMERSSFEK
NPIDFLEAKGYKEVKKDLIIKLPKYSLFELENGRKRMLASAKQLQKGNELALPSKYVNFLYLASH
YEKLKGSPEDNEQKQLFVEQHKHYLDEIIEQISEFSKRVILADANLDKVLSAYNKHRDKPIREQA
ENIIHLFTLTRLGAPRAFKYFDTTIDPKQYRSTKEVLDATLIHQSITGLYETRIDLSQLGGDSGGS
GGSGGSTNLSDIIEKETGKQLVIQESILMLPEEVEEVIGNKPESDILVHTAYDESTDENVMLLTSD
APEYKPWALVIQDSNGENKIKMLSGGSKRTADGSEFEPKKKRKV*

EGFP-KASH

MVSKGEELFTGVVPILVELDGDVNGHKFSVSGEGEGDATYGKLTLKFICTTGKLPVPWPTLVTT
LTYGVQCFSRYPDHMKQHDFFKSAMPEGYVQERTIFFKDDGNYKTRAEVKFEGDTLVNRIELK
GIDFKEDGNILGHKLEYNYNSHNVYIMADKQKNGIKVNFKIRHNIEDGSVQLADHYQQNTPIGD
GPVLLPDNHYLSTQSALSKDPNEKRDHMVLLEFVTAAGITLGMDELYKSGLRSREEEEETDSR
MPHLDSPGSSQPRRSFLSRVIRAALPLQLLLLLLLLLACLLPASEDDYSCTQANNFARSFYPMLR
YTNGPPPT*

**Table S2**. Oligonucleotide Sequences Used for Base Editing of Splice Sites

| Base Editor | Gene | Exon | Sequence | PAM | Figure |
|---|---|---|---|---|---|
| ABE | TRAP1 | 4 | GTAGGTGTTTATACGGGAGC | TGA | 1F-G |
| ABE | TRAP1 | 4 | TGTAGGTGTTTATACGGGAG | CTG | 1F-G |
| ABE | TRAP1 | 4 | CTGTAGGTGTTTATACGGGA | GCT | 1B-C, 1F-G |
| ABE | TRAP1 | 4 | CCTGTAGGTGTTTATACGGG | AGC | 1F-G |
| ABE | TRAP1 | 4 | TCCTGTAGGTGTTTATACGG | GAG | 1B-C, 1F-G |
| ABE | TRAP1 | 4 | TTCCTGTAGGTGTTTATACG | GGA | 1B-C, 1F-G |
| ABE | RAB34 | 9 | GTCAGGTGAGAATGTCCGAG | AAT | 1B-C, 1F-G |
| ABE | RAB34 | 9 | TGTCAGGTGAGAATGTCCGA | GAA | 1B-C, 1F-G |
| ABE | RAB34 | 9 | GTGTCAGGTGAGAATGTCCG | AGA | 1B-C, 1F-G |
| ABE | RAB34 | 9 | TGTGTCAGGTGAGAATGTCC | GAG | 1B-C, 1F-G |
| ABE | Col4A5 | 21 | ACTCAGGTGATGAGATATGT | GAA | 1B-C |
| ABE | Col4A5 | 21 | TACTCAGGTGATGAGATATG | TGA | 1B-C |
| ABE | Col4A5 | 21 | TCTTACTCAGGTGATGAGAT | ATG | 1B-C |
| ABE | APP | 17 | TTTTCAAGGTGTTCTTTGCA | GAA | 1B-C, 4B-C, E, G-H, K, 5B, S3F, S4B |
| ABE | RELA | 7 | ACTGCCTCAGGTGCCCCCAA | CAC | S1B |
| ABE | RELA | 7 | TCACTGCCTCAGGTGCCCCC | AAC | 2C, |
| ABE | RELA | 7 | TCACTGCCTCAGGTGCCCCC | AAC | 2C, S1B |
| ABE | RELA | 7 | TATACCTTTCTGCACCTTGT | CAC | S1B |
| ABE | JAG1 | 12 | TTCTAGGATTTGGTTAATGG | TTA | 2B |
| ABE | JAG1 | 12 | TCACCTGACAGAGGTTTCCA | GAG | 2B |
| ABE | HSF1 | 6 | CGCAGCTGTTCAGCCCCTCG | GTG | 2C |
| ABE | HSF1 | 6 | CCGCAGCTGTTCAGCCCCTC | GGT | S1B |
| ABE | HSF1 | 6 | TACGCACACTGGCCAGGCTG | CTG | 2C, S1B |
| ABE | HSF1 | 6 | CTACGCACACTGGCCAGGCT | GCT | S1B |
| ABE | LMNA | 11 | TCCCAGGGCTCCCACTGCAG | CAG | S1B |
| ABE | LMNA | 11 | TTCCCAGGGCTCCCACTGCA | GCA | 2B |
| ABE | LMNA | 11 | GACAACTCACCTGGGTTCGG | GGG | 2B,S1B |
| ABE | LMNA | 11 | AGACAACTCACCTGGGTTCG | GGG | S1B |
| ABE, SD 1 | APP | 17 | AAGTTTACCTACCTCCACCA | CAC | 4D,L,5C, S3F, S4C |
| ABE, SD 2 | APP | 17 | ACCTACCTCCACCACACCAT | GAT | 4D,L,5C |
| ABE, SD 3 | APP | 17 | AGTTTACCTACCTCCACCAC | ACC | 4D,L,5C |
| CBE | JAG1 | 12 | CTAGAAGAGGAGAAGGGGAG | AGA | 1B-C |
| CBE | JAG1 | 12 | TCCTAGAAGAGGAGAAGGGG | AGA | 1D-E, H-I, 2B |
| CBE | JAG1 | 12 | ATCCTAGAAGAGGAGAAGGG | GAG | 1D-E |
| CBE | JAG1 | 12 | AATCCTAGAAGAGGAGAAGG | GGA | 1D-E, H-I |
| CBE | JAG1 | 12 | AAATCCTAGAAGAGGAGAAG | GGG | 1D-E, H-I |
| CBE | JAG1 | 12 | ACCATTAACCAAATCCTAGAAG | AGG | 1D-E, H-I |
| CBE | JAG1 | 12 | CAAATCCTAGAAGAGGAGAA | GGG | 1D-E, H-I |
| CBE | RELA | 7 | CACCTGAGGCAGTGAAAACA | AGG | 1H-I |
| CBE | RELA | 7 | GCACCTGAGGCAGTGAAAAC | AAG | 1H-I |
| CBE | BAP1 | 2 | GAGGCCTGGGTGGGCGACA | AGA | 1H-I, 2D, S1B |
| CBE | BAP1 | 2 | CTTACCGAAATCTTCCACGA | GCA | 2D |

| | | | | | |
|---|---|---|---|---|---|
| CBE | BAP1 | 2 | CTCTTACCGAAATCTTCCAC | GAG | S1B |
| CBE | AHCY | 9 | CGGTCCACCTACACGCAGGC | AGG | 2D |
| CBE | AHCY | 9 | CCCACCTTCTTGGGCAGGAA | ATG | 2D |
| CBE | EGFR | 23 | ACCCCTGAGAGGATGAAGCA | AGA | 2B, S1B |
| CBE | EGFR | 23 | CACACTTGACCATGATCATG | TAG | 2B |
| CBE | EGFR | 23 | ACTCACACTTGACCATGATC | ATG | S1B |
| CBE | JAG1 | 12 | TTGCCACCACTCACCTGACA | GAG | 2B |
| CBE | APP | 17 | ACACCTTGAAAACAAATTAA | GAA | 4B |
| ABE | APP | SA OT 1 | TTTTAAAGGTGTTCTTTGCG | GAG | S3F |
| ABE | APP | SA OT 2 | TTTTGAAGGTGTTCTTTACA | GAG | S3F |
| ABE | APP | SA OT 3 | TTTACCAGGTGTTCTTTGCT | GAA | S3F |
| ABE | APP | SA OT 4 | TTATCAAGGTGTTCTATACA | GAA | S3F |
| ABE | APP | SA OT 5 | CTTTCAAGTTGTTCTGTGCA | GAG | S3F |
| ABE | APP | SD OT 1 | AAGTTTCCCTACCGCCACCA | TGG | S3F |
| ABE | APP | SD OT 2 | AAGGTTACCTTCCTCCACCA | TTC | S3F |
| ABE | APP | SD OT 3 | AAGTTTCCCTACATCCACCA | TTG | S3F |
| ABE | APP | SD OT 4 | AAGTCTACGTACCTCCACCA | TCT | S3F |
| ABE | APP | SD OT 5 | TAGTTTACCTCCCTCCACCC | TCT | S3F |

**Table S3**. Primer Sequences for DNA/RNA amplification and Sequencing

| Target | Exon | Purpose | Sequence |
|---|---|---|---|
| TRAP1 | 4 | gDNA FW | AACGATCACTTGAGCCTGGG |
| TRAP1 | 4 | gDNA RV | TGAGGGAGGTACCTGGATGG |
| TRAP1 | 4 | gDNA sequencing | GCAACATAGCGAGACCCTGT |
| RAB34 | 9 | gDNA FW | GGCTCTGCTGTGACACTCTT |
| RAB34 | 9 | gDNA RV | AGTAGGGGGCATCCAGTCTT |
| RAB34 | 9 | gDNA sequencing | AGCTCTGGAGGAATGAGGGT |
| EGFR | 23 | gDNA FW | TTGGGTCCTTACAGCAATCC |
| EGFR | 23 | gDNA RV | GATGCAAAGGCCTCAGCTGTTT |
| EGFR | 23 | gDNA sequencing | GAGGTAGACTGAGGCTTCCAGC |
| EGFR | 23 | RT-PCR FW | GCTCAACTGGTGTGTGCAGATC |
| EGFR | 23 | RT-PCR RV | TCACGGAACTTTGGGCGACTAT |
| EGFR | 23 | NGS cDNA FW | TCGTCGGCAGCGTCAGATGTGTATAAGAGACAG GCTCAACTGGTGTGTGCAGATC |
| EGFR | 23 | NGS cDNA RV | GTCTCGTGGGCTCGGAGATGTGTATAAGAGACAG TCACGGAACTTTGGGCGACTAT |
| JAG1 | 12 | gDNA FW | GCTGAAGCCCTGTGTTTGTG |
| JAG1 | 12 | gDNA RV | GCTGTAGGGACTGCCAATGT |
| JAG1 | 12 | gDNA sequencing | TCGTCAGTATCTCCTTCCACC |
| JAG1 | 12 | RT-PCR FW | CGGATTTAAGTGTGTGTGCCCC |
| JAG1 | 12 | RT-PCR RV | TTCTGGCAGGGATTAGGCTCAC |
| JAG1 | 12 | NGS cDNA FW | TCGTCGGCAGCGTCAGATGTGTATAAGAGACAGCGGATTTAAGTGT GTGTGCCCC |
| JAG1 | 12 | NGS cDNA RV | GTCTCGTGGGCTCGGAGATGTGTATAAGAGACAGTTCTGGCAGGGA TTAGGCTCAC |
| BAP1 | 2 | gDNA FW | TGAATAAGGGCTGGCTGGAG |
| BAP1 | 2 | gDNA RV | ATTGTGTGACCGGGGTCTTC |
| BAP1 | 2 | gDNA sequencing | GTAGGGTTCCTGGCACTGTC |
| BAP1 | 2 | RT-PCR FW | GTCTGTGTGTGGGACTGAGG |
| BAP1 | 2 | RT-PCR RV | GTTGGGTATCAGCTGGTGGG |
| BAP1 | 2 | NGS cDNA FW | TCGTCGGCAGCGTCAGATGTGTATAAGAGACAG GTCTGTGTGTGGGACTGAGG |
| BAP1 | 2 | NGS cDNA RV | GTCTCGTGGGCTCGGAGATGTGTATAAGAGACAG GTTGGGTATCAGCTGGTGGG |
| AHCY | 9 | gDNA FW | AGAGACGGGCTTTCACTGTG |
| AHCY | 9 | gDNA RV | TTAGACACGTGACCCTTGGC |
| AHCY | 9 | gDNA sequencing | CGGCCAGCTCAACATTTCTT |
| AHCY | 9 | RT-PCR FW | CATCTTTGTCACCACCACAGGC |
| AHCY | 9 | RT-PCR RV | AGGTACTGGGCTTGCTTCTCAG |
| AHCY | 9 | NGS cDNA FW | TCGTCGGCAGCGTCAGATGTGTATAAGAGACAG GTCAAGTGGCTCAACGAGAACG |
| AHCY | 9 | NGS cDNA RV | GTCTCGTGGGCTCGGAGATGTGTATAAGAGACAG TCCAAGACCACTGAGCTCATGG |
| RELA | 7 | gDNA FW | CAGATTGGCACCCACTGGACTA |
| RELA | 7 | gDNA RV | CCTGGTCCCGTGAAATACACCT |

| | | | |
|---|---|---|---|
| RELA | 7 | gDNA sequencing | GGGTCAGTGTGTCTAACCCTCC |
| RELA | 7 | RT-PCR FW | CCCACGAGCTTGTAGGAAAGGA |
| RELA | 7 | RT-PCR RV | CCTGGTCCCGTGAAATACACCT |
| RELA | 7 | NGS cDNA FW | TCGTCGGCAGCGTCAGATGTGTATAAGAGACAG CCCACGAGCTTGTAGGAAAGGA |
| RELA | 7 | NGS cDNA RV | GTCTCGTGGGCTCGGAGATGTGTATAAGAGACAG CCTGGTCCCGTGAAATACACCT |
| LMNA | 11 | gDNA FW | GTCCCCAAGTCTTCCTGAGC |
| LMNA | 11 | gDNA RV | TCCTACCCCTCGATGACCAG |
| LMNA | 11 | gDNA sequencing | TGTCTTCCCTCTCCTCCTCC |
| LMNA | 11 | gDNA sequencing | ACCCATCTCCTCTGGCTCTT |
| LMNA | 11 | RT-PCR FW | GACGACGAGGATGAGGATGGAG |
| LMNA | 11 | RT-PCR RV | CACCCCTTTCCCTTGGCTTCTA |
| LMNA | 11 | NGS cDNA FW | TCGTCGGCAGCGTCAGATGTGTATAAGAGACAG ACGGCTCTCATCAACTCCAC |
| LMNA | 11 | NGS cDNA RV | GTCTCGTGGGCTCGGAGATGTGTATAAGAGACAG TGAGGAGGACGCAGGAAG |
| HSF1 | 11 | gDNA FW | CTGTTCTGACTTCCCTCCCTCC |
| HSF1 | 11 | gDNA RV | TGGGACTTGGCTCACCTGAATC |
| HSF1 | 11 | gDNA sequencing | TGGGACTTGGCTCACCTGAATC |
| HSF1 | 11 | RT-PCR FW | TGCCTGGACAAGAATGAGCTCA |
| HSF1 | 11 | RT-PCR RV | CTCTAGGAGACAGTGGGGTCCT |
| HSF1 | 11 | NGS cDNA FW | TCGTCGGCAGCGTCAGATGTGTATAAGAGACAG TGCCTGGACAAGAATGAGCTCA |
| HSF1 | 11 | NGS cDNA RV | GTCTCGTGGGCTCGGAGATGTGTATAAGAGACAG CTCTAGGAGACAGTGGGGTCCT |
| APP | 17 | gDNA FW | CCAACCAGTTGGGCAGAGAATA |
| APP | 17 | gDNA RV | ACCTGGTTGTATTGAGAGGCCA |
| APP | 17 | gDNA sequencing | AGTTCTTAGCAAAAAGCTAAGCCTAA |
| APP | 17 | RT-PCR FW | GAAGTTGAGCCTGTTGATGCCC |
| APP | 17 | RT-PCR RV | CGTTCTGCTGCATCTTGGACAG |
| APP | 17 | NGS cDNA FW | TCGTCGGCAGCGTCAGATGTGTATAAGAGACAGGAAGTTGAGCCTG TTGATGCCC |
| APP | 17 | NGS cDNA RV | GTCTCGTGGGCTCGGAGATGTGTATAAGAGACAGCGTTCTGCTGCA TCTTGGACAG |
| APP | 17 | NGS gDNA FW (mice experiments) | TCGTCGGCAGCGTCAGATGTGTATAAGAGACAG ATCCAAATGTCCCCTGCAT |
| APP | 17 | NGS gDNA RV (mice experiments) | GTCTCGTGGGCTCGGAGATGTGTATAAGAGACAG CATGGAAGCACACTGATTCG |
| APP | 17 | NGS cDNA FW (mice experiments) | TCGTCGGCAGCGTCAGATGTGTATAAGAGACAG ACACAGAAAACGAAGTTGAGC |
| APP | 17 | NGS cDNA RV (mice experiments) | GTCTCGTGGGCTCGGAGATGTGTATAAGAGACAG GCGATAATGAGTAAATCATAAA |
| APP | SA OT 1: hg38 chr8: 9530933 6- | NGS gDNA FW | TCGTCGGCAGCGTCAGATGTGTATAAGAGACAG TTTGACCAGATCCCTCTCTCA |

| | | | |
|---|---|---|---|
| | 95309936 | | |
| APP | SA OT 1: hg38 chr8: 95309336-95309936 | NGS gDNA RV | GTCTCGTGGGCTCGGAGATGTGTATAAGAGACAGGGGTCTGGTCCCTACCCTTA |
| APP | SA OT 2: hg38 chr1: 58551430-58551930 | NGS gDNA FW | TCGTCGGCAGCGTCAGATGTGTATAAGAGACAGCAACCTTCCAACCAAGCAAT |
| APP | SA OT 2: hg38 chr1: 58551430-58551930 | NGS gDNA RV | GTCTCGTGGGCTCGGAGATGTGTATAAGAGACAGCAGTGTGACCCTTCAAGCAA |
| APP | SA OT 3: hg38 chr7: 81700000-81700496 | NGS gDNA FW | TCGTCGGCAGCGTCAGATGTGTATAAGAGACAGACAGCCAAACCAAAAATTGC |
| APP | SA OT 3: hg38 chr7: 81700000-81700497 | NGS gDNA RV | GTCTCGTGGGCTCGGAGATGTGTATAAGAGACAGTGGGTGGAATTGACAATGAA |
| APP | SA OT 4: hg38 chr8: 107193000-107193476 | NGS gDNA FW | TCGTCGGCAGCGTCAGATGTGTATAAGAGACAGGTGCATGGGATGTTCATCAG |
| APP | SA OT 4: hg38 chr8: 107193000-107193477 | NGS gDNA RV | GTCTCGTGGGCTCGGAGATGTGTATAAGAGACAGCTCAGCCAGACTGCCAACTT |
| APP | SA OT 5: hg38 chr4: 94419456-94419956 | NGS gDNA FW | TCGTCGGCAGCGTCAGATGTGTATAAGAGACAGTGCAAGGCCCAAGTAATAGC |

| | | | |
|---|---|---|---|
| APP | SA OT 5: hg38 chr4: 94419456-94419957 | NGS gDNA RV | GTCTCGTGGGCTCGGAGATGTGTATAAGAGACAG CATTTTCAGCACCAGAACCA |
| APP | SD OT 1: hg38 chr2: 238138000-238138478 | NGS gDNA FW | TCGTCGGCAGCGTCAGATGTGTATAAGAGACAG CCTGTTTCTTAGCCGTCCTG |
| APP | SD OT 1: hg38 chr2: 238138000-238138479 | NGS gDNA RV | GTCTCGTGGGCTCGGAGATGTGTATAAGAGACAG CACGTGGGAAAACGTGATTA |
| APP | SD OT 2: hg38 chr13: 80179531-80179999 | NGS gDNA FW | TCGTCGGCAGCGTCAGATGTGTATAAGAGACAG CCCCAAGATATGGGATGTTG |
| APP | SD OT 2: hg38 chr13: 80179531-80179999 | NGS gDNA RV | GTCTCGTGGGCTCGGAGATGTGTATAAGAGACAG TTAGCAAGCCCCTTTTGTTC |
| APP | SD OT 3: hg38 chr16: 35480069-35480569 | NGS gDNA FW | TCGTCGGCAGCGTCAGATGTGTATAAGAGACAG ATGGACTTGGTCCCAAACAG |
| APP | SD OT 3: hg38 chr16: 35480069-35480569 | NGS gDNA RV | GTCTCGTGGGCTCGGAGATGTGTATAAGAGACAG AGCTCCACCATATCGTGTCC |
| APP | SD OT 4: hg38 chr9: 88814241-88814741 | NGS gDNA FW | TCGTCGGCAGCGTCAGATGTGTATAAGAGACAG CGGCTGGCCATACACTTTTA |
| APP | SD OT 4: hg38 | NGS gDNA RV | GTCTCGTGGGCTCGGAGATGTGTATAAGAGACAG TGCCTGAGATGTGAGTTTGC |

| | | | |
|---|---|---|---|
| | chr9: 88814241-88814742 | | |
| APP | SD OT 5: hg38 chr10: 16760535-16760999 | NGS gDNA FW | TCGTCGGCAGCGTCAGATGTGTATAAGAGACAG<br>CTGGGACATGAAGCTCGTCT |
| APP | SD OT 5: hg38 chr10: 16760535-16761000 | NGS gDNA RV | GTCTCGTGGGCTCGGAGATGTGTATAAGAGACAG<br>GCACTCCAGCCTAGGTGACA |
| APP | | qPCR | GAGGTACCCACTGATGGTAATG |
| APP | | qPCR | CTGAATCCCACTTCCCATTCT |
| GAPDH | | qPCR | CAATGACCCCTTCATTGACC |
| GAPDH | | qPCR | TTGATTTTGGAGGGATCTCG |
